# Optimal Filters for ERP Research I: A General Approach for Selecting Filter Settings

**DOI:** 10.1101/2023.05.25.542359

**Authors:** Guanghui Zhang, David R. Garrett, Steven J. Luck

## Abstract

Filtering plays an essential role in event-related potential (ERP) research, but filter settings are usually chosen on the basis of historical precedent, lab lore, or informal analyses. This reflects, in part, the lack of a well-reasoned, easily implemented method for identifying the optimal filter settings for a given type of ERP data. To fill this gap, we developed an approach that involves finding the filter settings that maximize the signal-to-noise ratio for a specific amplitude score (or minimizes the noise for a latency score) while minimizing waveform distortion. The signal is estimated by obtaining the amplitude score from the grand average ERP waveform (usually a difference waveform). The noise is estimated using the standardized measurement error of the single-subject scores. Waveform distortion is estimated by passing noise-free simulated data through the filters. This approach allows researchers to determine the most appropriate filter settings for their specific scoring methods, experimental designs, subject populations, recording setups, and scientific questions. We have provided a set of tools in ERPLAB Toolbox to make it easy for researchers to implement this approach with their own data.

## 1. Introduction

Filters are essential in research using event-related potentials (ERPs). At a minimum, a low-pass filter must be applied in hardware prior to digitizing the continuous EEG signal so that aliasing can be avoided (Picton et al., 2000). Low-pass filters can also minimize muscle artifacts and induced electrical noise (de Cheveigné & Nelken, 2019; Luck, 2014), and high-pass filters can significantly improve statistical power by reducing skin potentials (Kappenman & Luck, 2010). However, a strong filter may reduce the amplitude of the ERP component of interest as well as attenuating noise. Moreover, inappropriate filter settings can create temporal smearing, artifactual deflections, or bogus oscillations in the ERP waveform, potentially leading to erroneous conclusions (Acunzo et al., 2012; de Cheveigné & Nelken, 2019; Rousselet, 2012; Tanner et al., 2015; Vanrullen, 2011; Widmann et al., 2015; Yeung et al., 2007).

What, then, are the ideal filter settings for a given study? Although some recommendations have been proposed (e.g., Duncan et al., 2009; Luck, 2014; Widmann et al., 2015), the existing cognitive and affective ERP literature does not provide a clear, complete, and quantitatively justified answer to this question. This is partly because different filter settings may be optimal for different experimental paradigms, participant populations, scoring methods, and scientific questions. As a result, the range of filter settings used in published ERP research varies widely across laboratories and often within a laboratory, presumably based on a combination of empirical testing, mathematical understanding (or misunderstanding) of filtering, and lab lore.

The goal of the present paper is to provide a principled and straightforward approach for determining optimal filter settings. The approach is conceptually simple: A set of filters are evaluated, and the optimal filter is the one that maximizes the data quality without producing unacceptable levels of waveform distortion. However, this requires methods for quantifying the data quality for a specific amplitude or latency score and methods for assessing waveform distortion. Our approach includes methods for each of these steps^1^.

A companion paper (Zhang et al., 2023) uses this approach to provide recommendations for seven commonly-used ERP components combined with four different scoring methods (mean amplitude, peak amplitude, peak latency, and 50% area latency). That paper uses data from the ERP CORE (Compendium of Open Resources and Experiments; Kappenman et al., 2021), which includes data from 40 young adults who performed six standardized paradigms that yielded seven commonly-studied ERP components.

However, the optimal filter settings for those data may not be optimal for different populations of participants (e.g., infants), different recording setups (e.g., dry electrodes), or different ERP components (e.g., the contingent negative variation). For example, data quality for the error-related negativity is much poorer in young children (Isbell & Grammer, 2022) than in adults (Kappenman et al., 2021), and this may necessitate different filter settings. The present paper therefore describes in detail how researchers can apply this approach to their own data. To make this approach easy to implement, we have also added new tools to version 9.2 and higher of ERPLAB Toolbox (Lopez-Calderon & Luck, 2014). Our approach can be implemented either through ERPLAB’s graphical user interface or through scripting, and we have also provided example scripts at https://osf.io/98kqp/.

Our intention is for this paper to be useful for researchers from a broad variety of backgrounds, whether or not they have any technical expertise with signal processing. Consequently, the technical details have been deemphasized in the main text and can be found in footnotes, supplementary materials, or the cited papers.

## 2. Review of basic filter properties

We first review the basic frequency-domain properties of the kinds of filters used most often in ERP research (see Widmann et al., 2015, for a more detailed overview). Figure 1 shows the *frequency response functions* of several different filters along with their effects on an example averaged ERP waveform. Figure 1a shows low-pass filters (which pass low frequencies and attenuate high frequencies), and Figure 1b shows high-pass filters (which pass high frequencies and attenuate low frequencies). In the ERP waveforms, the low-pass filters mainly appear to “smooth” the waveform, whereas the high-pass filters mainly appear to reduce the amplitude of the P3 wave.

**Figure 1.**
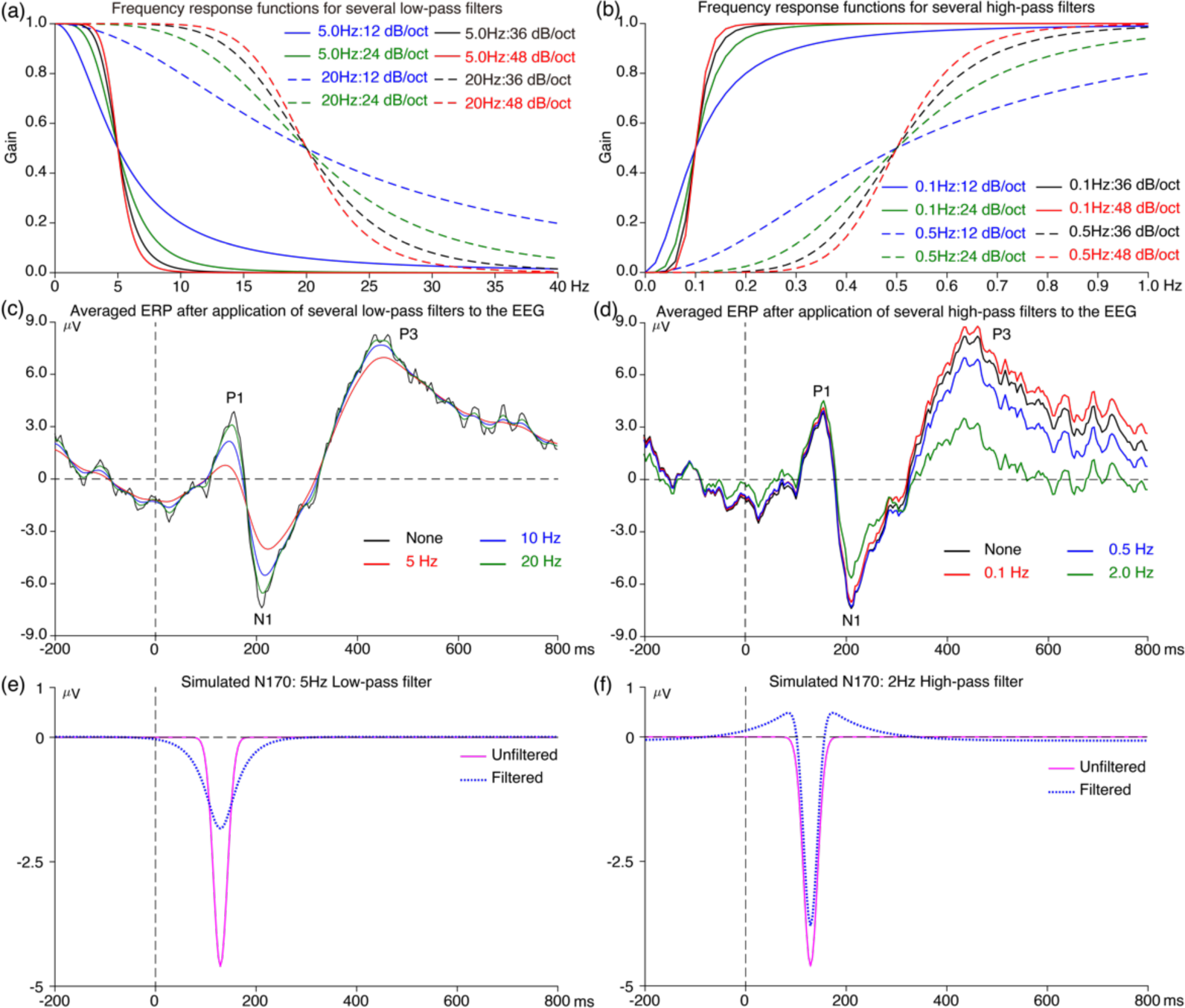
Frequency response functions for several filters and their application to a single-participant ERP waveform and to a simulated N170 waveform. The frequency response functions quantify the extent to which a given frequency is passed versus blocked by the filter. (a) Frequency response functions for low-pass filters. Two cutoff frequencies are shown (5 Hz and 20 Hz), combined with four roll-off slopes (12 dB/octave, 24 dB/octave, 36 dB/octave, and 48 dB/octave). (b) Frequency response functions for high-pass filters. Two cutoff frequencies are shown (0.1 Hz and 0.5 Hz), combined with four roll-off slopes (12 dB/octave, 24 dB/octave, 36 dB/octave, and 48 dB/octave). (c) Averaged ERP waveform with different half-amplitude low-pass filter cutoffs (no filter, 5 Hz, 10 Hz, and 20 Hz, all with slopes of 12 dB/octave). (d) The same averaged ERP waveform with different half-amplitude high-pass filter cutoffs (no filter, 0.1 Hz, 0.5 Hz, and 2 Hz, all with slopes of 12 dB/octave). (e) and (f) Simulated N170 waveform filtered by a 5Hz low-pass filter and 2Hz high-pass filter (12 dB/octave). Note that the filters used for (c) and (d) were applied to the continuous EEG data prior to epoching and averaging. All filters used here were noncausal Butterworth filters, and cutoff frequencies indicate the half-amplitude point. The waveforms in (c) and (d) were from the face condition in the ERP CORE N170 paradigm, Subject 40, CPz electrode site.

The frequency response functions characterize how the filters impact the frequency content of the data. The horizontal axis is frequency, and the vertical axis is gain. The gain indicates the extent to which a given frequency passes through the filter rather than being attenuated. A gain of 1.0 means that the frequency passes through completely (no filtering); a gain of 0.75 means that 75% of the signal at that frequency passes through the filter and that the signal is attenuated by 25% at that frequency; a gain of 0.0 means that the signal is completely attenuated at that frequency. The frequency response function of a filter is often summarized with the *half-amplitude cutoff frequency*, which is the frequency at which the gain is 0.5 and the signal is attenuated by 50%. The high-pass filters shown in Figure 1a have a half-amplitude cutoff at either 0.1 Hz or 0.5 Hz. The low-pass filters shown in Figure 1b have a half-amplitude cutoff at either 5 Hz or 20 Hz. When both high-pass and low-pass filters are applied, their combined effects are referred to as the *bandpass* of the filter (e.g., a bandpass of 0.1–5Hz for the combine effects of a 0.1 Hz high-pass filter and a 5 Hz low-pass filter).

Filters are also characterized by their roll-offs, which specify how rapidly the gain changes as the frequency changes. This is often summarized by the slope of the filter at its steepest point, using logarithmic units of decibels of gain change per octave of frequency change (dB/octave). The filters in Figure 1 have roll-offs ranging from relatively shallow (12 dB/octave) to relatively steep (48 dB/octave). Researchers often assume that a steeper roll-off is better, because this means that most frequencies are either passed nearly completely (a gain near 1.0) or attenuated nearly completely (a gain near 0.0). As we will demonstrate, however, steeper roll-offs tend to produce greater waveshape distortion.

Filters can be either *causal* (unidirectional) or *noncausal* (bidirectional). Causal filters create a rightward shift in the waveform and are typically avoided except under specific conditions (Rousselet, 2012; Woldorff, 1993). All filters used here were therefore noncausal.

Although filters can be extremely valuable in attenuating noise, they inevitably distort the time course of the ERP waveform (see Luck, 2014; Widmann et al., 2015). Low-pass filters typically “smear” the waveform, causing ERP components to begin artificially early and end artificially late. This is illustrated for a simulated N170 waveform in Figure 1e. When a 5 Hz lowpass filter was applied to the simulated waveform, the onset of the component was shifted earlier in time and the offset was shifted later in time. By contrast, high-pass filters typically produce artifactual opposite-polarity deflections before and after an ERP component. For example, Figure 1f shows that applying a 2 Hz high-pass filter produced artifactual positive peaks before and after the simulated N170 component. Thus, it is important to consider the time-domain waveform distortion produced by a filter as well as the filter’s frequency-domain properties.

## 3. Overview of the present approach

In this section, we provide a brief overview of our approach to determining optimal filter parameters. The following sections will provide a more detailed explanation and justification. We assume that the component of interest in a given study can be isolated from other ERP components by means of a difference wave; Section 8.3 will discuss how our approach can be modified if this assumption is not valid.

Figure 2 provides a flowchart for our approach when the score of interest is an amplitude score (e.g., N170 peak amplitude). Section 7 will discuss how the procedure is modified for latency scores (e.g., N170 peak latency). The approach consists of two parallel streams of analysis, one examining the effects of a set of candidate filters on data quality and another examining the effects of the candidate filters on waveform distortion. For amplitude measures, data quality is quantified using a special version of the *signal-to-noise ratio* (SNR), and waveform distortion is quantified using the relative size of the artifactual peaks produced by the filter (the *artifactual peak percentage* or APP). An SNR value and an APP value are obtained for each candidate filter, and the filter that produces the best SNR without exceeding a criterion for waveform distortion is selected as the optimal filter. Note that this general procedure could be used with a different metric of data quality, a different metric of waveform distortion, or a different threshold for waveform distortion.

**Figure 2.**
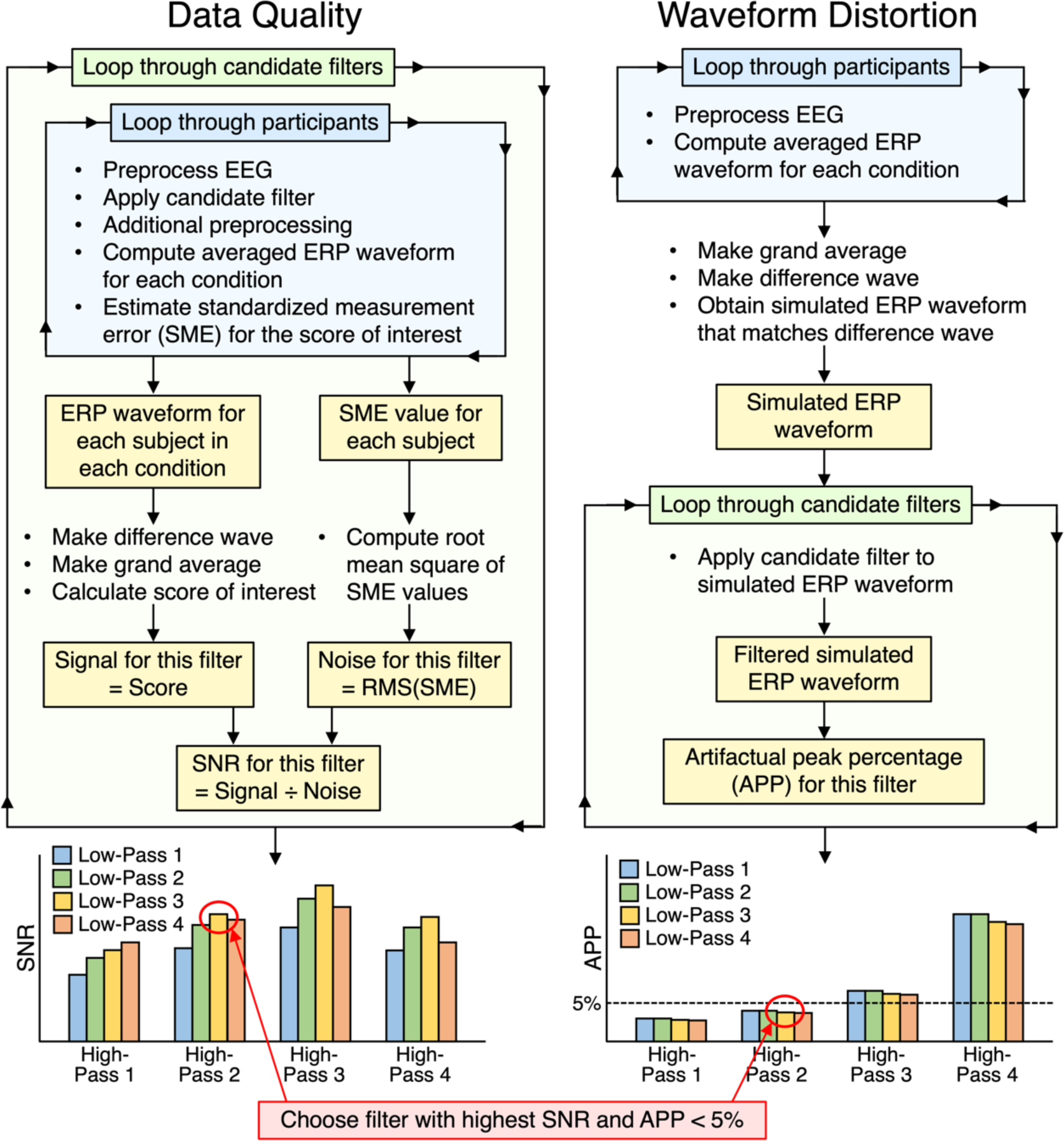
Flow chart of procedure for selecting the optimal filter for an ERP amplitude score. For latency scores, the noise is used instead of the signal-to-noise ratio, and the filter with the lowest RMS(SME) is chosen from among those that yield an artifactual peak percentage of less than 5%. If there is reason to believe that the grand average difference waveform is not a good reflection of the true effect, the right side of the flowchart can be repeated with multiple different simulated ERP waveforms that reflect the range of possibilities.

The set of candidate filters can be chosen by examining the range of filters used in prior research. Alternatively, it can be chosen by starting with a broad set of combinations of low-pass and high-pass filters, determining the general range of parameters that yield good results, and then repeating the with a finer set of combinations within that range.

Ordinarily, one set of data should be used to select the optimal filter parameters, and then those parameters should be applied to new data. If the optimal filters are selected on the basis of one set of data are then applied to the same data, this “double dipping” will likely increase the false positive rate. There may be exceptions, but a careful justification would be needed before applying the parameters to the same dataset that was used to select the parameters. The previous studies used to determine the filter parameters need not be identical to the study of interest: reasonable filtering parameters can be chosen as long as prior recordings are available that contain reasonably similar waveforms and noise levels. We have also added options in ERPLAB’s Channel Operations tools that allow users to add noise of various types to prior data.

## 4. Defining and quantifying the signal-to-noise ratio

This section describes our approach to estimating the data quality resulting from a given set of filter parameters (the left side of Figure 2). We use the signal-to-noise ratio as our metric of data quality because filters can decrease the size of the signal as well as reduce the noise, and it is important to determine whether the reduction in noise outweighs the loss of signal. Although the concept of SNR has been used in ERP research for many decades, it is not simple to define and quantify either the signal or the noise in a way that is truly useful. This section therefore provides a new way of visualizing, defining, and quantifying the signal, the noise, and the SNR.

The signal portion of the SNR is usually framed in terms of the amplitude of the signal, so we will focus on ERP amplitudes for the next several sections. A slightly different approach is required for ERP latencies, as described in Section 7.

### 4.1. An informal visualization of the effects of filters on signal and noise

The logic behind applying high-pass and low-pass filters to EEG/ERP data is that some types of noise are confined to relatively low frequencies (e.g., skin potentials) whereas other types of noise are confined to relatively high frequencies (e.g., line noise and muscle artifacts). The signal of interest (i.e., the neural activity) typically has less power than the noise in the very low frequencies and/or the very high frequencies, so attenuating these frequencies may reduce the noise more than it reduces the signal. This subsection provides visualizations of how filtering reduces both the signal and the noise.

To illustrate the loss of signal, Figure 1c shows the effects of low-pass filters with different cutoffs^2^ on an averaged ERP waveform. These filters “smoothed” out the high-frequency noise, but reducing the low-pass cutoff frequency also reduced the P1 and N1 amplitudes. These low-pass filters had less impact on the amplitude of the P3 wave. In general, low-pass filters will decrease the amplitude of fast, narrow waves such as P1 and N1, with less impact on slow, broad waves such as P3 and N400.

Figure 1d shows the effects of high-pass filtering on the same averaged ERP waveform. P3 amplitude declined as the cutoff frequency increased. However, P1 and N1 amplitude were largely unaffected until the cutoff reached 2.0 Hz. In general, high-pass filters will decrease the amplitude of the longer-latency, broader waves but will have less impact on shorter-latency, narrower waves.

Although low-pass filters produce a clearly visible reduction in high-frequency noise in the averaged ERP waveforms (as in Figure 1c), the noise reduction produced by high-pass filters is not usually obvious in the averaged ERP waveform (as in Figure 1d). The effects of high-pass filtering can be more easily visualized by looking at the single-trial EEG epochs, as illustrated in Figure 3. Slow drifts in the EEG (Figure 3a) cause the single-trial EEG waveforms to tilt upward on some trials and downward on other trials. Because the epochs are baseline-corrected using the prestimulus period, the trial-to-trial variability in voltage increases progressively over the course of the epoch. In the corresponding averaged ERP waveform (Figure 3b), this leads to a progressive increase in the standard error of the mean over the course of the epoch. Note that the participant shown in Figure 3 had unusually large low-frequency drifts, which makes the slow voltage drifts easier to see. However, virtually all participants exhibit increasing drift and an increasing standard error of the mean over the course of the epoch if the prestimulus interval is used for baseline correction. These low-frequency drifts add uncontrolled variability to the averaged ERP waveforms, and this can cause a dramatic reduction in statistical power, especially for later components that are farther away from the baseline period (Acunzo et al., 2012; Hennighausen et al., 1993; Kappenman & Luck, 2010).

**Figure 3.**
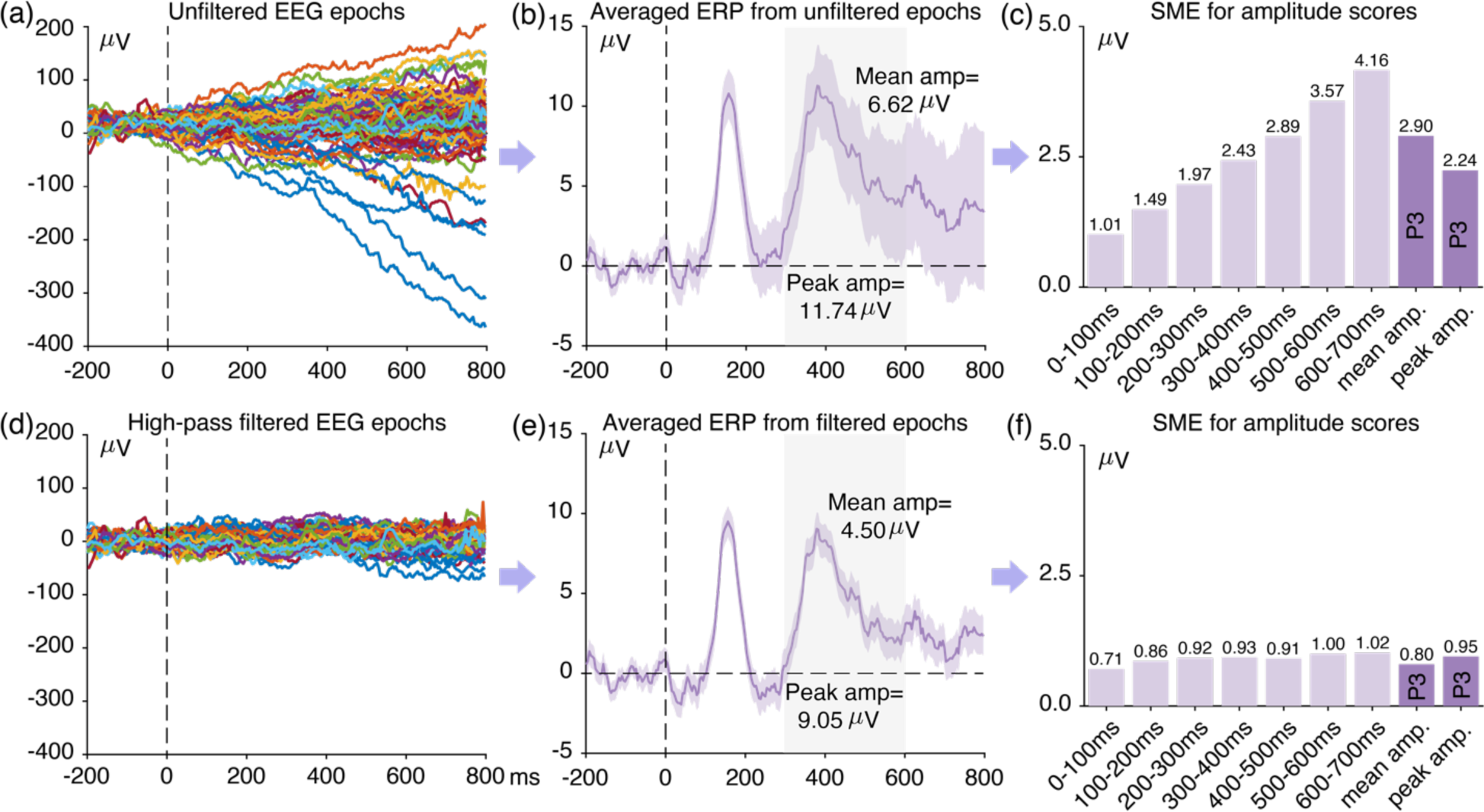
Examples of single-trial EEG epochs, averaged ERP waveforms, and standardized measurement error (SME) values without filtering (top row) and after application of a high-pass filter with a half-amplitude cutoff at 0.5 Hz (bottom row). The SME was calculated for the mean amplitude in consecutive 100-ms time periods, for mean amplitude in the P3 measurement window (300-600 ms), and for peak amplitude in the P3 measurement window. The shaded region for the ERP waveforms reflects the standard error of the mean at each individual time point. The filter was a noncausal Butterworth filter with a slope of 12 dB/octave. The data were from the standard condition in the ERP CORE visual oddball paradigm, Subject 40, Pz electrode site.

Drift in the single-trial EEG can be dramatically reduced by applying a 0.5 Hz high-pass filter to the EEG (Figure 3d), and this also reduces the standard error of the mean, especially later in the epoch (Figure 3e). Thus, low-frequency drift is a threat to statistical power but can be minimized by high-pass filtering.

### 4.2. The classic definition of signal-to-noise ratio (SNR) and its limitations

Now that we have seen how filters can influence both the signal and the noise, we will consider how to quantify the signal-to-noise ratio (SNR). Classically, the SNR in ERP research is defined separately at each individual time point in the averaged ERP waveform. The signal is the amplitude of the averaged waveform at a given time point, and the noise is quantified by some measure of trial-to-trial variability at that time point (Picton et al., 2000). Each point in an averaged ERP waveform is the mean of the single-trial voltages at that time point, so the standard error of the mean is a natural way to quantify the noise. Thus, the SNR at a given time point can be quantified as the amplitude at that time point (the signal) divided by the standard error of the mean at that time point (the noise). The standard error of the mean (SEM) is typically estimated using Equation 1:

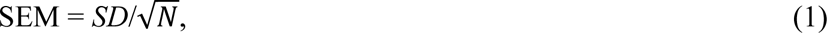

where *SD* is the standard deviation of the single-trial amplitudes at that time point and *N* is the number of trials being averaged together. Because the standard error of the mean is linearly related to 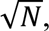 this definition of the SNR explains why the SNR of an averaged ERP waveform improves linearly with 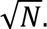

In Figures 3b and 3e, we could compute the SNR at each time point by dividing the amplitude of the averaged ERP at that time point by the standard error of the mean at that time point. However, the SNR at individual time points is not particularly useful in most ERP research, because most studies derive amplitude and latency *scores* from the pattern of voltages across multiple time points. For example, the size of mismatch negativity (MMN) component in the ERP CORE (Kappenman et al., 2021) was measured as the mean voltage from 125–225 ms in the deviant-minus-standard difference waves. Because this is the type of score that is typically entered into statistical analyses and used to test the hypotheses of a study, the SNR of a given score is much more important than the SNR at individual time points (except when mass univariate analyses are used). However, there is no simple mathematical relationship between the SNR of a score that is derived from multiple time points and the SNR at the individual time points.

In addition, the effect of a given filter will depend on what scoring method is being used (e.g., mean amplitude vs. peak amplitude). For example, when the mean voltage over a reasonably wide time window is used to score the amplitude of an ERP component, high-frequency noise has relatively little impact on the score. This is illustrated in Figure 4, which shows a simulated ERP waveform with and without high-frequency noise. The rapid upward and downward noise deflections within the measurement window tend to cancel each other out, and the mean amplitude from 300-500 ms in the noisy waveform is similar to the mean amplitude from 300-500 ms in the clean waveform. However, the peak amplitude is strongly affected by the high-frequency noise. High-frequency noise will also have a substantial impact on peak latency scores. By contrast, low-frequency noise has a modest impact on peak latency scores but a large impact on mean amplitude and peak amplitude scores. Thus, there is no such thing as a generic metric of noise; the noise must be defined with respect to the specific method used to score the amplitude or latency. When choosing a filter, it is therefore essential to consider how the averaged ERP waveform will be scored and how the filter will impact the SNR of that specific score.

**Figure 4.**
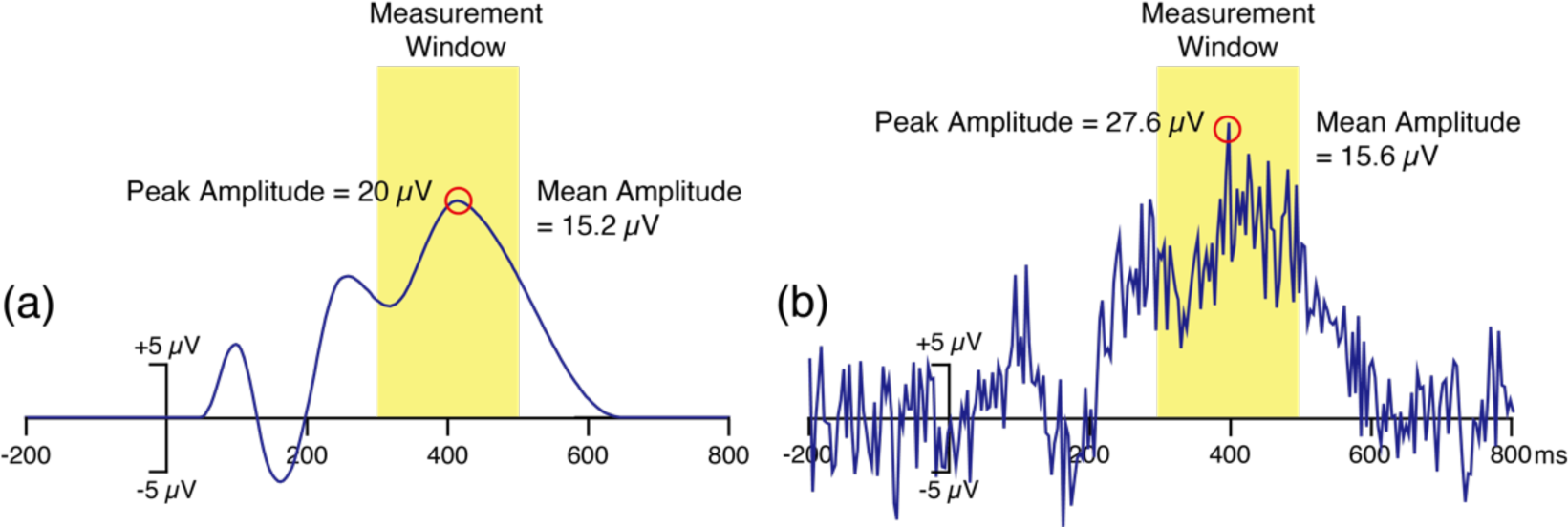
Simulated ERP waveform without noise (a) and with high-frequency noise added (b). This high-frequency noise had very little impact on the mean amplitude during this window because the upward and downward noise deflections largely canceled each other. However, the noise had a large impact on the peak amplitude.

A partial solution to this problem is to define the signal as the mean voltage during the time window of interest and the noise as the standard deviation of the voltage across the points during the baseline period (e.g., Debener et al., 2008; Klug & Gramann, 2021). However, this is relevant only for mean amplitude scores and does not apply to other scoring methods (e.g., peak amplitude, peak latency). In addition, there is no guarantee that the noise level will remain constant between the baseline period and the time window of interest. The noise might increase owing to low-frequency drifts (as in Figure 3a), or it might decrease owing to stimulus-induced suppression of alpha-band activity (Klimesch, 2012). In addition, it is difficult to estimate the effects of low-frequency noise in the prestimulus interval, where low-frequency drifts are minimized by the baseline correction procedure. Indeed, we have found that the baseline noise level often provides a misleading estimate of the noise that impacts a given amplitude or latency score (see, e.g., Figure S4 in Luck et al., 2021). Thus, we need a means of quantifying the SNR that can apply to any scoring method and that directly reflects the noise that impacts the score of interest.

### 4.3. Using the standardized measurement error (SME) to estimate the SNR for ERP amplitude scores

A new definition of SNR that meets these criteria was recently proposed by Luck et al. (2021). The *signal* is straightforward: It can be estimated by the score itself (although a caveat will be described in Section 4.4). For example, when MMN peak amplitude is measured from a deviant-minus-standard difference wave, the signal is the measured peak amplitude. The *noise* can then be estimated as the *standard error of measurement* for that score (which reflects both the trial-to-trial variability and the number of trials being averaged together).

This is a simple generalization of the method for computing the SNR at each individual time point, in which the standard error of the mean at a given time point was used to estimate the noise. The value at a given time point in an averaged ERP waveform is the mean of the single-trial voltages at that time point. Because it is a mean, the standard error of *measurement* for this value is the standard error of the *mean*. Thus, the SNR at a given time point is the mean across trials at that time point divided by the standard error of the mean at that time point.

However, when the signal of interest is a score that is based on the pattern of voltages across multiple time points, we need to estimate the standard error of measurement for that particular score. Luck et al. (2021) developed an approach for quantifying the standard error of measurement for ERP amplitude and latency scores, and the resulting estimate of the noise is called the *standardized measurement error* (SME; see Section S5 of the supplementary materials for a conceptual overview). Thus, the SNR for a given amplitude score can be quantified as the score divided by the SME for that score. We refer to this specific definition of SNR as *SNR_SME_*. As one would expect, SNR_SME_ depends on the size of the score (the signal) along with the amount of trial-to-trial variability and the number of trials (which are combined together in the SME).

Equation 1 can be used to estimate the SME when the amplitude of an ERP component is scored as the mean voltage within a given time window in an averaged ERP waveform (Luck et al., 2021). That is, the mean voltage across the time period is scored for each individual trial, and Equation 1 is applied to these values. When estimated using this simple analytic approach, the result is called the *analytic SME* or *aSME*. Unfortunately, this simple approach is not valid for other scoring methods, such as peak amplitude. For those scoring methods, Luck et al. (2021) developed a bootstrapping method for estimating the SME. The result is called the *bootstrapped SME* or *bSME*. Our bootstrapping procedure is described in Section S6 of the supplementary materials. Note that the SME assumes that the score of interest will be obtained from averaged ERPs, not from single trials. Some other yet-to-be-developed approach would be needed when single-trial scores are used as the dependent variable in statistical analyses.

The aSME is automatically computed by version 8.1 and later of ERPLAB Toolbox. Computing the bSME currently requires Matlab scripting, but the scripts are relatively simple, and example scripts are available at https://doi.org/10.18115/D58G91. In addition, the ERP CORE resource contains SME values and the code required to compute them for all seven ERP components (https://doi.org/10.18115/D5JW4R; see Zhang & Luck, 2023).

No matter how the SME is computed, it is an estimate of the standard error of measurement for the score of interest, and it can therefore be used as the noise term when computing the SNR_SME_. For example, P3 amplitude in the ERP CORE was scored as the mean amplitude from 300-600 ms in the target-minus-standard difference waves, and the SNR_SME_ for P3 amplitude is this score divided by the SME of the score.

Using this approach, the SNR_SME_ can be estimated for both filtered and unfiltered data to determine the extent to which a given filter increases or decreases the signal-to-noise ratio. This is illustrated in the rightmost column of Figure 3. When P3 amplitude was scored as the mean amplitude from 300-600 ms in the unfiltered data, the score was 6.62 µV (see Figure 3b) and the SME of this score was 2.90 µV (see Figure 3c). The SNR_SME_ was therefore 6.62/2.90 or 2.28. When the peak amplitude was scored instead, the score was 11.74 µV and the SME of this score was 2.24 µV, yielding an SNR_SME_ of 11.74/2.24 or 5.24. After a 0.5 Hz high-pass filter was applied (Figure 3d-f), the mean amplitude and peak amplitude scores were slightly smaller than before (4.50 µV and 9.05 µV, respectively). However, the SME values were reduced by a much greater amount (to 0.80 µV and 0.95 µV, respectively). Consequently, the SNR_SME_ was almost doubled by the filtering to 5.63 for mean amplitude and 9.53 for peak amplitude.

This example shows how we can determine which filter parameters lead to the best signal-to-noise ratio. To our knowledge, this is the first method that can quantify how filters impact the SNR of the actual amplitude scores that are used to test hypotheses in most cognitive and affective ERP experiments.

As shown in the flowchart in Figure 2, the SNR_SME_ can be computed for each candidate filter to determine which filter yields that best data quality. The following sections provide some important details about how the signal is defined and how single-participant SNR_SME_ values should be aggregated across conditions and across participants. In addition, it is important to keep in mind that the SNR is not the only factor that should be considered when choosing a filter. In particular, Section 6 will show that a filter with a better SNR_SME_ may produce more waveform distortion than a filter with a worse SNR_SME_.

### 4.4. Improving the estimate of the signal

Although it is straightforward to use the amplitude score as the signal in the SNR_SME_ calculation, these scores are distorted by any noise in the averaged ERP waveform and are therefore an imperfect estimate of the signal. For example, Figure 4 shows that high-frequency noise will cause the peak amplitude to be overestimated, which will then lead to an overestimate of the SNR_SME_. Filtering out the high-frequency noise will decrease the peak amplitude, bringing it closer to the true value, but this might create the illusion that filtering has decreased the signal-to-noise ratio.

A simple solution to this problem is to obtain the score from the grand average ERP waveform, which typically has much less noise than the single-participant waveforms. This score could then be divided by the SME for a given participant to estimate the SNR_SME_ for that participant. To obtain a group SNR_SME_, the score from the grand average would be divided by an aggregate of the single-participant SME values (see Section 4.7).

Obtaining the score from the grand average is not a perfect solution, because some noise will remain in the grand average and contribute to the estimate of the signal. This residual noise is often negligible, but when substantial noise remains in the grand average an artificial ERP waveform can instead be used to estimate the signal (see Section 5). We found nearly identical results for the ERP CORE data when measuring the signal from the grand average or from artificial waveforms, so we used the grand average when calculating the SNR_SME_ in the present paper and in the companion paper (Zhang et al., 2023).

Obtaining the score from the grand average is also an imperfect approach for nonlinear scoring methods, such as peak amplitude, because the mean of the single-participant peaks is not the same as the peak of the grand average waveform. For example, if the timing of an ERP component varies across participants, the peak amplitude of the grand average ERP waveform will be smaller than the average of the single-participant peaks (even in the absence of noise). However, the goal of the present procedure is not to determine the true SNR_SME_, but instead to determine how the SNR_SME_ varies across different filter settings. The pattern of SNR_SME_ values across filters is typically not impacted by the nonlinearity problem, so measuring from the grand average typically works well in practice for determining the optimal filtering parameters.

### 4.5. Measuring from difference waves

An averaged ERP waveform is the weighted sum of many underlying components that overlap in time and space (Nunez & Srinivasan, 2006). To isolate a specific ERP component, many experiments focus on differences between experimental conditions (e.g., oddballs versus standards for P3 and MMN, faces versus cars for N170). In these cases, we recommend estimating both the signal and the SME from difference waves (e.g., oddballs-minus-standards, faces-minus-cars). The reasoning is illustrated in Figure 5, which shows the grand average ERP waveforms from the ERP CORE N2pc experiment (Kappenman et al., 2021). The N2pc component is defined as the difference between the waveform at electrode sites contralateral versus ipsilateral to the target location (indicated by yellow shading). Although this difference is approximately 1.5 µV, the N2pc is superimposed on a broad positivity arising from other ERP components, bringing the overall voltage up to approximately 5 µV. If we measured the signal from the contralateral and ipsilateral waveforms (the *parent* waveforms), we would get a value that is approximately three times as large as the actual 1.5 µV N2pc component. This would vastly overestimate the size of the signal. In addition, if we measured from the parent waveforms, a high-pass filter that reduced the broad positivity might appear to reduce the signal even if it had minimal impact on N2pc amplitude. Similarly, if we estimated the SME from the parent waveforms, our measure of the noise would also be distorted by the overlapping components.

**Figure 5.**
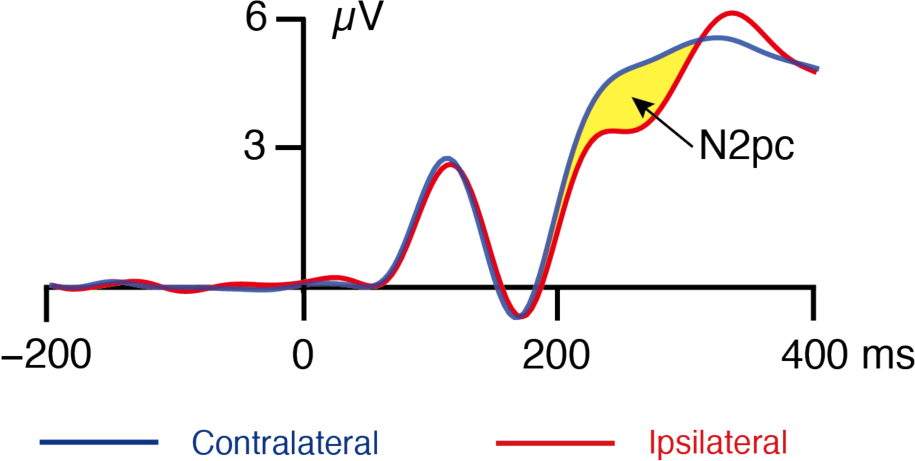
Grand average ERP waveforms from the ERP Core N2pc experiment. Separate waveforms are shown for trials where the target was contralateral to the electrode site and trials where the target was ipsilateral to the electrode site. The N2pc is defined as the difference in voltage between the contralateral and ipsilateral waveforms (denoted here by yellow shading).

By measuring both the signal and the SME from the difference waveform, we can avoid these problems and more directly determine how different filters impact the signal of interest and the noise that impacts that signal. Note that the signal is measured from the difference wave of the grand average across participants, whereas the noise is measured from each individual participant and then aggregated across participants (see Figure 2).

It is important to note that difference waves do not always perfectly isolate a single component. In addition, there may be cases in which difference waves are not available or would actually mischaracterize the effect of interest. We consider these issues in Section 8.

### 4.6. Obtaining the SME of a score obtained from a difference wave

When an amplitude or latency score is obtained from a difference wave, it is ordinarily necessary to use bootstrapping to estimate the SME. However, when the score of interest is the mean amplitude over a fixed time window, it is faster and easier to obtain the analytic SME values provided by ERPLAB for the individual conditions and use the following equation to estimate the SME corresponding to the difference wave:

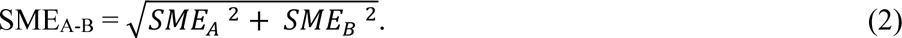

In this equation, *SME_A-B_* is the SME of the difference between conditions A and B, and *SME_A_* and *SME_B_* are the SMEs of the two individual conditions. Note that this equation applies only when the score is the mean voltage across a time window. In addition, it applies only when the difference wave is between waveforms from separate trials (e.g., target minus standard for P3, unrelated minus related for N400), not when it is a difference between two electrode sites (e.g., contralateral minus ipsilateral for N2pc or lateralized readiness potential). Additional mathematical details are provided in Section S8 of the supplementary materials.

### 4.7. Computing an SNR_SME_ value that reflects the entire sample participants

Up to this point, we have discussed how to use the SME to estimate the noise level for each individual participant. However, we need a way to aggregate these values across participants to estimate the overall noise level for a given set of filter parameters. This could be accomplished by simply averaging the single-participant SME values. However, participants with particularly noisy data have an outsized effect on statistical power, and it is better to use the *root mean square* (*RMS*) of the single-participant SME values as the noise estimate for the group (Luck et al., 2021). The RMS is obtained by squaring each single-participant SME value, taking the mean of these squared values, and then taking the square root of this mean:

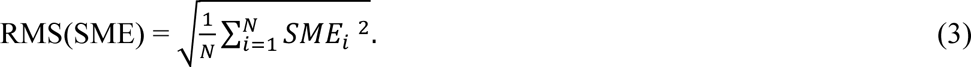

In this equation, SME_i_ is the SME value for participant *i*, of the *N* participants after combining across conditions using Equation 2.

The resulting RMS(SME) value provides an aggregate estimate of the noise level for a given score after the application of a given set of filter parameters. As shown in Figure 2, the SNR_SME_ value for that set of parameters is then computed by dividing the signal (the score obtained from the filtered grand average difference wave) by the RMS(SME) value.

## 5. Assessing waveform distortion

This section describes our approach to estimating the waveform distortion produced by a given set of filter parameters (the right side of Figure 2). The most straightforward way to assess time-domain filter distortion is to pass an artificial waveform through the filter and compare the filtered and unfiltered versions of this waveform. Artificial waveforms must be used for this purpose because the true (i.e., noise-free) waveform is not usually known for real data, making it difficult to know if the filter is “revealing” the true waveform by eliminating noise or is instead creating a bogus effect that mischaracterizes the underlying brain activity (Yeung et al., 2007).

To create an appropriate artificial waveform, it is necessary to estimate the waveshape of the real ERP effect. In many cases, the grand average difference wave (e.g., faces minus cars for the N170 component) provides a good starting point. This difference wave can be used to create a noise-free artificial waveform with the key properties of the experimental effect. If there is reason to believe that the grand average difference wave is not a good reflection of the actual waveshape, the shape of the artificial waveform can be systematically varied, and the effects of filtering can be assessed across a range of waveshapes. Section 8 describes strategies that can be applied if multiple components are present in the difference wave.

Note that the grand average waveform used as a starting point for creating the simulated ERP waveform should be created using minimal filtering. Otherwise, it may already contain significant filter distortions.

### 5.1. Creating simulated N170 and P3 effects

This subsection provides examples of artificial waveforms that simulate two effects from the ERP CORE: a) the larger N170 for faces than for cars in a visual discrimination paradigm, and b) the larger P3 for oddballs than for standards in a visual oddball paradigm. We have chosen these two effects because they span the gamut from a relatively early perceptual effect to a relatively late cognitive effect. These and most other ERP effects can be simulated with Gaussian and ex-Gaussian functions^3^. Beginning in version 9.2, ERPLAB Toolbox provides a tool for using these and other functions to create artificial waveforms that simulate ERP components. Whereas this tool simulates averaged ERP waveforms, the SEREEGA toolbox (Krol et al., 2018) can be used to simulate single-trial EEG epochs that contain ERP-like effects.

Panels a and b of Figure 6 show the grand average N170 and P3 effects from the ERP CORE. The N170 paradigm involved a series of faces, cars, scrambled faces, and scrambled cars, and the N170 effect was defined as the faces-minus-cars difference. The P3 paradigm involved a sequence of target (rare) and standard (frequent) letter categories, and the P3 effect was defined as the target-minus-standard difference. The N170 effect can be approximated by a negative-going Gaussian function with a mean of 129 ms and an SD of 14 ms. The P3 waveform is typically skewed to the right, and the ERP CORE P3 effect can be approximated by an ex-Gaussian function with a Gaussian mean of 310 ms, a Gaussian SD of 58 ms, and an exponential rate parameter (*λ*) of 2000 ms.

**Figure 6.**
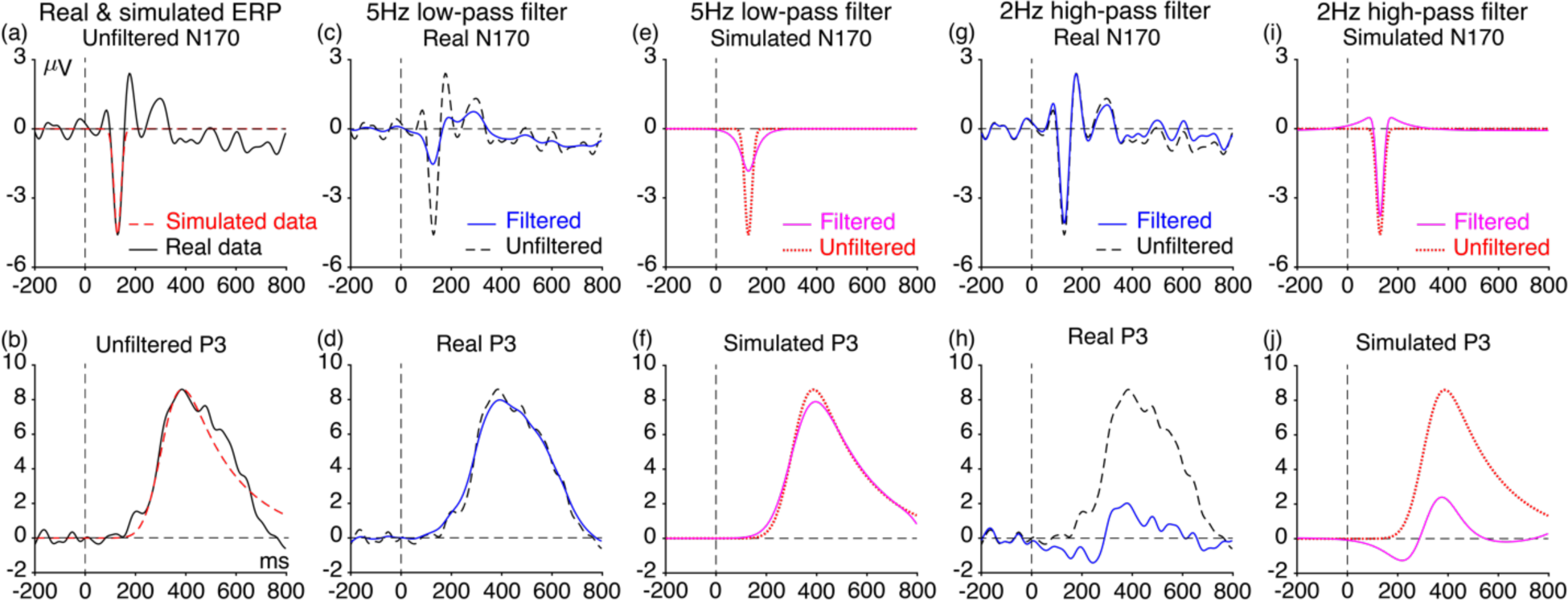
Filter-induced distortions of real and simulated N170 and P3 components from the ERP CORE. (a) Grand average N170 difference wave and simulation with a Gaussian function (mean = 129 ms, SD = 14 ms, peak amplitude = -4.6 *µV*). (b) Grand average P3 difference wave and simulation with an ex-Gaussian function (mean = 310 ms, SD = 58 ms, *λ = 2000ms*, peak amplitude *=8.6 µV*). The artificial waveforms were preceded and followed by 1000 ms of zero values to avoid edge artifacts. (c, d) Effects of a 5 Hz low-pass filter on the real N170 and P3 waveforms, respectively. (e, f) Effects of a 5 Hz low-pass filter on the simulated N170 and P3 waveforms, respectively. (g, h) Effects of a 2 Hz high-pass filter on the real N170 and P3 waveforms, respectively. (i, j) Effects of a 2 Hz high-pass filter on the simulated N170 and P3 waveforms, respectively. All filters used here were noncausal Butterworth filters with a slope of 12 dB/octave, and cutoff frequencies indicate the half-amplitude point.

These simulated waveforms are overlaid on the observed grand average waveforms in Figures 6a and 6b. They are not a perfect fit, but they do a reasonable job of capturing the key properties of the N170 and P3 components, and most ERP components can be approximated by Gaussian and ex-Gaussian functions with appropriate parameters.

Because the true waveform is not known, it may be necessary to create several different artificial waveforms that reflect different possibilities for the true waveform. A filter can then be chosen that minimizes the waveform distortion for the entire set of simulated waveforms. In addition, some effects may consist of changes in multiple overlapping components. This can be approximated by creating simulations of the individual components and then summing them together.

The following subsections show how the real and simulated waveforms shown in Figure 6 are distorted by a low-pass filter and a high-pass filter. We have chosen relatively extreme cutoff frequencies for these examples to make the distortions obvious. We also provide examples of the distortions produced by more typical filters in Figures 7 and 8.

**Figure 7.**
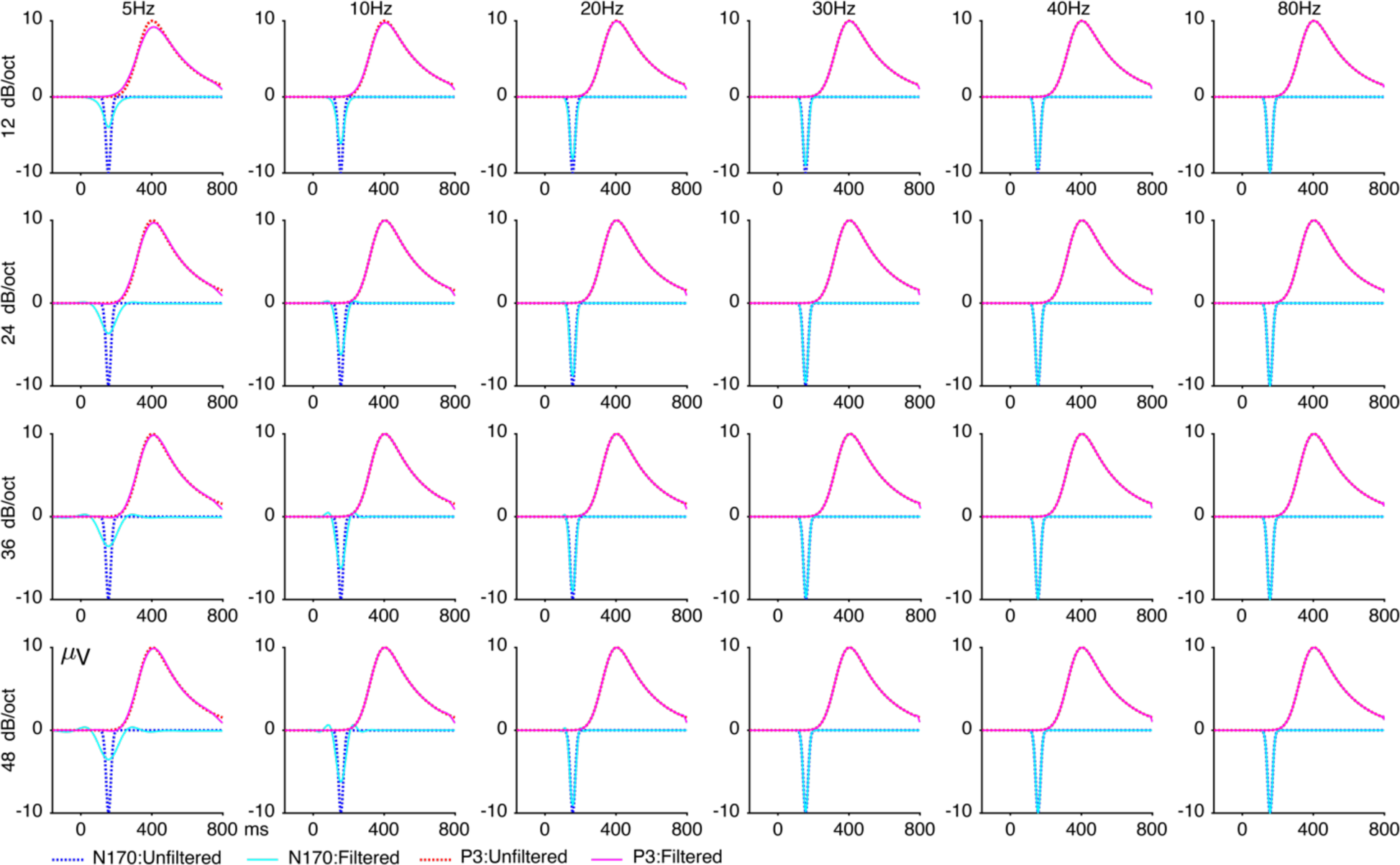
Effects of different low-pass filter cutoffs (5, 10, 20, 30, 40, and 80 Hz) and roll-offs (12, 24, 36, and 48 dB/octave) on the simulated N170 and P3 waveforms. Note that the distortion is most notable for the lowest cutoff frequencies. All filters used here were noncausal Butterworth filters with a slope of 12 dB/octave, and cutoff frequencies indicate the half-amplitude point.

**Figure 8.**
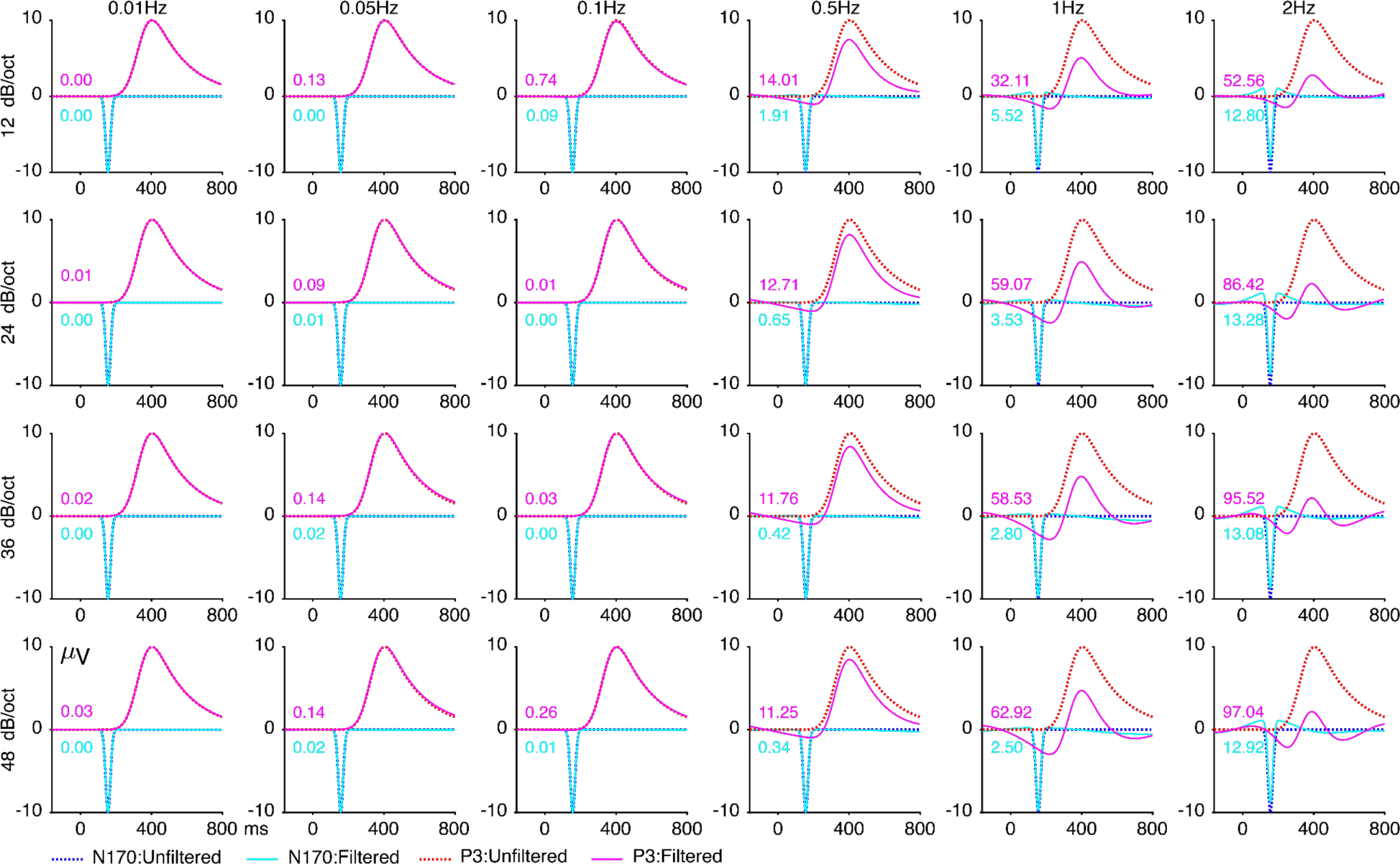
Effects of different high-pass filter cutoffs (0.01, 0.05, 0.1, 0.5, 1, and 2 Hz) and roll-offs (12, 24, 36, and 48 dB/octave) on the simulated N170 and P3 waveforms. Note that the distortion is most notable for the highest cutoff frequencies. All filters used here were noncausal Butterworth filters with a slope of 12 dB/octave, and cutoff frequencies indicate the half-amplitude point. The embedded number in each panel is the artificial peak percentage (shown in magenta for P3 and light blue for N170).

### 5.2. Effects of low-pass filtering on simulated ERPs

Panels c and d of Figure 6 show the results of applying a 5 Hz low-pass filter to the real N170 and P3 waveforms, and Panels e and f show the results of filtering the simulated versions of these waveforms. The filter reduced the amplitude of both the real and simulated N170 peaks. Figure 6e also shows that the filter “smeared out” the simulated N170, artificially creating an earlier onset time and a later offset time. By contrast, the 5 Hz low-pass filter had relatively little effect on the P3 wave (Figures 6d and 6f). Thus, as noted in Section 4.1, low-pass filters have a much larger effect on short-latency, narrow peaks such as the N170 than on long-latency, broad peaks such as the P3.

Figure 7 shows the effects of a variety of different low-pass filter cutoffs and roll-offs on the simulated N170 and P3 waveforms. When a relatively gentle roll-off of 12 dB/octave was used, the waveform distortion consisted of a progressively greater temporal smearing as the cutoff frequency declined, with minimal smearing when the cutoff was above 15 Hz. When steep roll-offs were used, however, the distortion of the simulated N170 also included opposite-polarity peaks on either side of the N170 (see, e.g., the cutoff of 10 Hz with a slope of 48 dB/octave). Thus, filtering N170 data with a steep slope might cause a researcher to reach the invalid conclusion that faces elicit a small, early, positive response as well as the typical N170 response. Filtering with a shallow slope avoids this problem. However, filters with shallow slopes still distort the onset and offset times of the waveform, especially with cutoff frequencies below 20 Hz. Whether these distortions are a significant problem depends on the nature of the scientific questions being asked and the analysis procedures being applied.

### 5.3. Effects of high-pass filtering on simulated ERPs

Panels g-j of Figure 6 show the results of applying a 2 Hz high-pass filter to real and simulated N170 and P3 waveforms. This filter did not produce any obvious distortion of the real N170 waveform, except for a modest reduction in peak amplitude, but the simulated waveform shows that the filter also produced opposite-polarity artificial peaks on each side of the simulated N170 wave. The other voltage deflections in the real data made it difficult to see these artifactual peaks. The filter produced much greater distortion of the P3 wave, dramatically reducing P3 amplitude and producing an artifactual negative peak prior to the true peak. The artifactual negative peak might lead to the incorrect conclusion that the target stimuli elicited a larger negativity than the standard stimuli (for additional examples of invalid conclusions that may arise from filtering, see Tanner et al., 2015; Yeung et al., 2007).

Figure 8 shows the effects of a variety of different high-pass filter cutoffs and roll-offs on the simulated N170 and P3 waveforms. The artifactual opposite-polarity peaks were minimal for cutoffs of 0.1 Hz or lower, but they became clearly visible at 0.5 Hz and increased progressively as the cutoff increased further. Note that the artifactual peaks were more pronounced prior to the P3 peak than after the P3 peak. This is a result of the right skew in the simulated P3 waveform. The same asymmetry can be observed in the filter artifacts for the real P3 waveform in Figure 6h. Thus, for right-skewed waveforms like the P3, high-pass filters produce a larger artifactual peak before than after the true peak.

### 5.4. Quantifying waveform distortion with the artifactual peak percentage (APP)

To quantify the waveform distortion produced by a given filter, we compute the *artifactual peak percentage* (APP). The APP reflects the amplitude of the artifactual peak produced by the filter relative to the amplitude at the peak of the true component after filtering. Absolute values are used so that a greater distortion always produces a larger value. Specifically, the APP is calculated from the filtered waveform as:

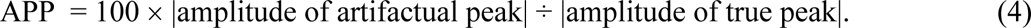

Consider, for example, the artificial P3 wave after high-pass filtering with a cutoff at 2 Hz and a slope of 12 dB/octave (Figure 8, upper right corner). The peak amplitude of the artifactual peak was -1.466 µV, and the peak amplitude of the true peak was +2.789 µV, so the artifactual peak percentage was 100 × |-1.466| ÷ |2.789| = 52.56%. This value is shown for each filter setting in Figure 8. Note that using peaks to quantify the size of a component can be problematic with real data, but it is not so problematic with noise-free artificial waveforms, and it has the advantage of simplicity. However, it would be reasonable for researchers to use an alternative measure, such as area amplitude, when computing the amplitude distortion percentage.

The idea behind this approach is that a small artifactual peak is likely to be obscured by the background noise and have no impact on the conclusions drawn from a given study, but a large artifactual peak might be statistically significant and lead to a bogus conclusion. In addition, the artifactual peak might be considered to be substantial in size if it is relatively large compared to the other peaks in the waveform (such as the true peak after filtering). For example, the artifactual peak in the upper right corner of Figure 8 looks like a very substantial effect when compared with the rest of the waveform.

Although artificial peaks are mainly a problem for high-pass filters, Figure 7 shows that they may also occur for low-pass filters with a steep cutoff (see, e.g., the lower left panel in Figure 7). We therefore provide the artifactual peak percentage values for the low-pass filters in supplementary Table S1.

### 5.5. Determining the maximal acceptable artifactual peak percentage (APP)

We define the optimal filter as the one that maximizes the data quality without producing unacceptable levels of waveform distortion. This requires setting a threshold for an unacceptably large APP.

Setting this threshold requires balancing the risk of a false positive (an artifactual peak that is large enough to be statistically significant) and the risk of a false negative (a true effect that is not statistically significant because of reduced SNR). This is analogous to the threshold for statistical significance in traditional frequentist statistical analyses (typically 0.05); a lower threshold such as 0.01 reduces the risk of a false positive (a Type I error) but also decreases statistical power and increases the risk of a false negative (a Type II error). Generally, scientists are more concerned about false positives than false negatives and choose a relatively conservative threshold. The chosen threshold is usually arbitrary, but at least the risks are well defined.

For most ERP studies conducted in low-noise environments with highly cooperative participant populations, we propose a threshold of 5% for the artifactual peak percentage. That is, we recommend that researchers use the filter parameters that produces the best SNR_SME_ while also producing an artifactual peak percentage of less than 5%. This amount of distortion would be like a 0.5 µV artifactual N2 preceding a 10 µV P3, a 0.4 µV artifactual P2 preceding an 8 µV N400, or a 0.1 µV artifactual P1 preceding a 2 µV N2pc. Artifactual effects of this size are unlikely to be statistically significant under typical conditions. If we increased the criterion to 10%, however, we might have a 1 µV artifactual N2, a 0.8 µV artifactual P2, or a 0.2 µV artifactual P1, which would have a good chance of being statistically significant^4^ and leading to a fundamentally incorrect conclusion. If we decreased the criterion to 1%, there would be almost no chance that the artifactual peaks would be significant, but we would also be choosing a filter that yields a poorer SNR and therefore lower statistical power. Thus, a maximal artifactual peak amplitude of 5% seems like a reasonable balance between false positives and false negatives.

However, we would like to stress that this 5% criterion is arbitrary, and it would not be straightforward to assess the actual probability that an artifactual effect of a given size would be statistically significant. Nonetheless, the 5% criterion seems reasonably conservative without being overly strict for most ERP studies conducted in low-noise environments with highly cooperative participant populations. Under other conditions, it is likely that a more liberal criterion would be justified. For example, in studies with noisy EEG signals or an unusually small number of trials, an artifactual peak of 10% amplitude is much less likely to lead to a statistically significant effect, and the boost in SNR and statistical power produced by a high-pass cutoff that produces an artifactual peak percentage of 10% may therefore be well justified.

When the amplitude of an ERP component is being scored, the temporal smearing of the waveform produced by a low-pass filter is not usually a concern (unless it impacts the SNR). Thus, we do not recommend using the amount of latency distortion as a criterion for filter section when mean or peak amplitudes are being scored. However, this smearing could be an issue when the exact onset or offset latency of an effect is of theoretical relevance or when mass univariate statistical analyses are used. In those situations, the specific theoretical questions should drive the decision about how much latency distortion is acceptable. Section S7 of the supplementary materials describes a latency distortion percentage metric that could be used for this purpose.

## 6. Example: Determining the optimal filter parameters for P3 mean amplitude

This section provides a concrete example of our approach, using data from the ERP CORE P3 paradigm (Kappenman et al., 2021) to select an optimal filter for P3 mean amplitude. This was an oddball paradigm with rare *targets* and frequent *standards*. The P3 was isolated using a target-minus-standard difference wave. We used candidate filters created by factorially combining high-pass cutoffs of 0, 0.01, 0.05, 0.1, 0.5, 1, and 2 Hz with low-pass cutoffs of 5, 10, 20, 30, 40, 80, and 115 Hz (12 dB/octave).

We will first describe the procedure for assessing the effects of data quality, which involves looping through each candidate filter (Figure 2, left). For each filter, we loop through each of the participants to obtain the single-participant averaged ERP waveforms and SME values for the targets and standards. This starts by applying the ordinary preprocessing steps that are needed for a given study. In the present case, these steps included shifting event codes to account for the monitor delay, downsampling the data to 256 Hz, re-referencing, and correcting for artifacts using independent component analysis (ICA). The candidate filter was then applied to the continuous data (to avoid edge artifacts). We then conducted additional preprocessing steps, which included epoching the data, performing baseline correction, and rejecting artifacts that were not corrected by ICA. A detailed description of these steps can be found in the companion paper (Zhang et al., 2023).

### 6.1. Quantifying the signal for each candidate filter

Averaged ERP waveforms for targets and for standard were obtained from the resulting data, along with the target-minus-standard difference wave. A grand average across participants was then computed for the difference wave. The score of interest for the P3 wave in the ERP CORE was the mean voltage from 300-600 ms at the Pz electrode site. This score was obtained from the grand average difference wave to serve as the estimate of the signal after the attenuation produced by the current candidate filter.

Figure 9a shows the resulting scores for each candidate (see supplementary Figure S1 for the grand average waveforms from which the scores were obtained and supplementary Figure S2 for the corresponding scalp maps). Significant attenuation of the P3 was produced by high-pass cutoffs above 0.1 Hz, whereas low-pass filtering had very little effect on the signal.

**Figure 9.**
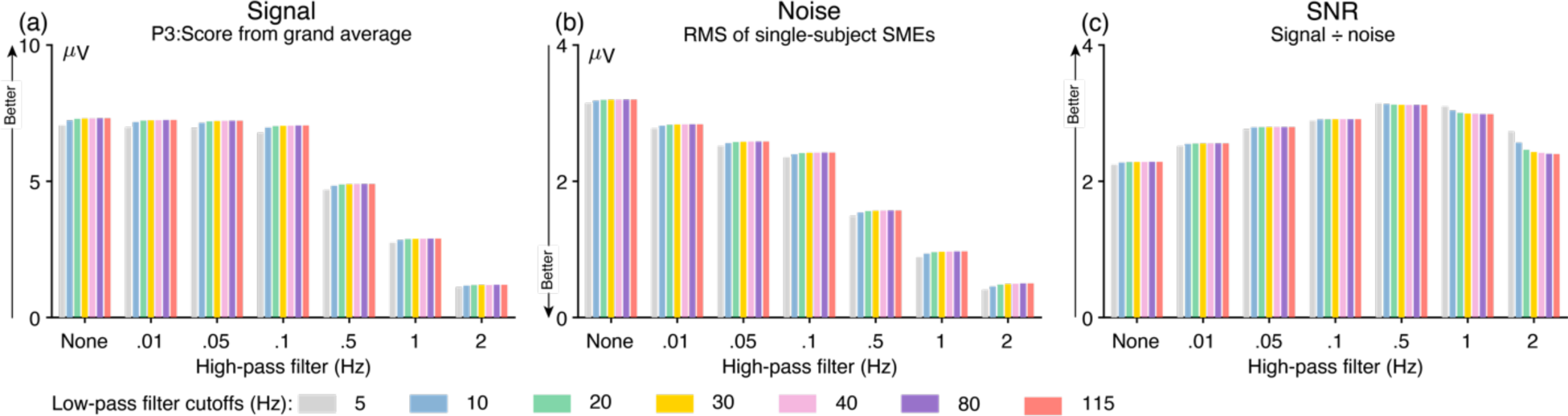
Demonstration of how the signal, noise, and signal-to-noise ratio (SNR) vary as a function of the filter settings for the P3 mean amplitude score from the ERP CORE visual oddball paradigm. (a) Signal: P3 mean amplitude score (from 300-600 ms at Pz) obtained from the grand average target-minus-standard difference wave. (b) Noise: Root mean square (RMS) of the single-participant standardized measurement error (SME) values for the P3 scores. Analytic SME values were obtained for each participant for the target and standard conditions for mean amplitude, and Equation 1 was applied to obtain the SME of the target-minus-standard difference for each participant. These SME values were then aggregated across participants using the RMS. (c) SNR: The signal divided by the noise for each filter setting. Note that filtering was applied to the continuous EEG prior to averaging with every combination of seven high-pass filter cutoffs (0, 0.01, 0.05, 0.1, 0.5, 1, and 2 Hz) and seven high-pass filter cutoffs (5, 10, 20, 30, 40, 80, and 115 Hz). All filters used here were noncausal Butterworth filters with a slope of 12 dB/octave, and cutoff frequencies indicate the half-amplitude point.

### 6.2. Quantifying the noise for each candidate filter

Because the score of interest was a mean amplitude value, ERPLAB was used to directly compute the SME for each condition using Equation 1. Bootstrapping would be needed to compute the SME for nonlinear measures, such as peak amplitude and peak latency.

We then applied Equation 2 to combine the SME values for the targets and the standards into a single SME value that reflects the data quality for the difference in amplitude between the targets and the standards. The SME values from the different participants were then aggregated using Equation 3 to obtain the RMS(SME), which reflects the data quality for the P3 amplitude score across the whole sample of participants.

Figure 9b shows the resulting RMS(SME) values for each of the candidate filters. The noise level decreased progressively as the high-pass cutoff increased. Thus, high-pass filtering decreased the signal (Figure 9a) and also decreased the noise (Figure 9b).

### 6.3. Quantifying the signal-to-noise ratio for each candidate filter

To assess whether the reduction in signal was outweighed by the reduction in noise for a given filter, we computed the SNR_SME_ for each candidate filter by taking the signal (the score from Figure 9a) and dividing it by the noise (the RMS(SME) from Figure 9b). The SNR_SME_ for each candidate filter is shown in Figure 9c.

The high-pass cutoff had a clear impact on the SNR_SME_, with the largest SNR_SME_ obtained at 0.5 Hz. Low-pass filtering had very little effect except when a 2 Hz high-pass filter was also applied. This is because mean amplitude scores are largely insensitive to high-frequency noise (see Figure 4). The best overall SNR_SME_ was obtained with a high-pass cutoff of 0.5 Hz and a low-pass cutoff of 5 Hz.

Supplementary Figure S2 shows how filtering impacts the scalp distribution of an ERP component and discusses how the SNR_SME_ might vary across scalp sites. Supplementary Figure S3 shows SNR_SME_ values for a denser sampling of high-pass cutoff frequencies between 0.1 and 1.0 Hz.

### 6.4. Assessing waveform distortion for each candidate filter

As shown in Figure 9c, the best SNR_SME_ for the P3 mean amplitude score was obtained with a high-pass cutoff of 0.5 Hz and no low-pass filtering. However, it is necessary to combine the information about SNR_SME_ with information about waveform distortion in order to select the optimal filter settings.

Our procedure for quantifying waveform distortion (Figure 2, right) begins by looping through the participants with the same processing steps used for the SME calculations, but with minimal filtering. In the present case, this led to an averaged ERP waveform for targets and for standards in each participant, which were used to create target-minus-standard difference waves. These were then combined across participants into a grand average difference wave. We then simulated a P3 difference wave to match the observed grand average difference wave (as shown in Figure 6b).

The next step was to loop through the candidate filters and quantify the amount of waveform distortion each filter produced in the simulated P3 waveform. Distortion was quantified with the artifactual peak percentage (APP). The top row of Figure 8 shows the filtered waveform and APP for each of the candidate high-pass cutoffs with no low-pass filtering, and supplementary Figure S3a shows the APP values for all the candidate filters.

As shown at the bottom of Figure 2, the final step of our procedure is to combine the data quality information with the waveform distortion information and select the filter with the best SNR_SME_ from among those with an APP of less than 5%. If we did not consider the APP, we would have chosen a high-pass cutoff at 0.5 Hz and a low-pass cutoff of 5 Hz, which produced the best SNR_SME_. However, Figure S3a shows this filter produced an APP of 12.2%, which was well above the 5% threshold. Lowering the high-pass cutoff to 0.1 Hz and keeping the low-pass cutoff at 5 Hz reduced the APP to 0.5%, well under the 5% threshold.

These results suggest that a bandpass of 0.1–5 Hz is optimal. However, after we determined that the best combination of data quality and waveform distortion lies somewhere between high-pass cutoffs of 0.1 and 0.5 Hz, we repeated the process with a denser sampling of cutoffs between 0.1 and 0.5 Hz (in steps of 0.1 Hz) to more precisely determine the optimal filter settings (see supplementary Figure S3). Among the filters with an APP of <5%, the best SNR_SME_ was produced with a high-pass cutoff of 0.2 Hz; there were no differences among the different low-pass cutoffs with this high-pass cutoff. Thus, when P3 mean amplitude is scored in studies like the ERP CORE visual oddball experiment, a high-pass cutoff of 0.2 Hz is optimal, along with whatever low-pass cutoff is useful given the other goals of the study. For example, a low-pass cutoff at 20 or 30 Hz can be applied to remove “fuzz” in plots of the waveforms that might otherwise make it difficult to visualize differences between conditions.

The companion paper provides recommendations for six other ERP components, and includes peak amplitude, peak latency, and 50% area latency scores in addition to mean amplitude scores. One could also assess the impact of different roll-off slopes, but a slope of 12 dB/octave is usually best in terms of minimizing waveform distortion (see Figures 7 and 8).

## 7. Selecting optimal filter settings for latency scores

The kinds of filters typically used in ERP research will typically decrease the amplitude of an ERP component (see Section 8 for exceptions). As a result, filters tend to reduce the difference in amplitude between groups or conditions. This is illustrated in Figures 10a and 10b, which show a simulation of two conditions in which the P3 amplitude differs. In the unfiltered data (Figure 10a), the peak amplitude differed by 0.5 µV between the two conditions. When a high-pass filter with a half-amplitude cutoff of 1 Hz was applied (Figure 10b), the difference in amplitude between conditions was reduced to 0.24 µV. This is why our approach to determining optimal filtering parameters for amplitude scores involves quantifying the effects of filtering on the signal as well as the noise.

**Figure 10.**
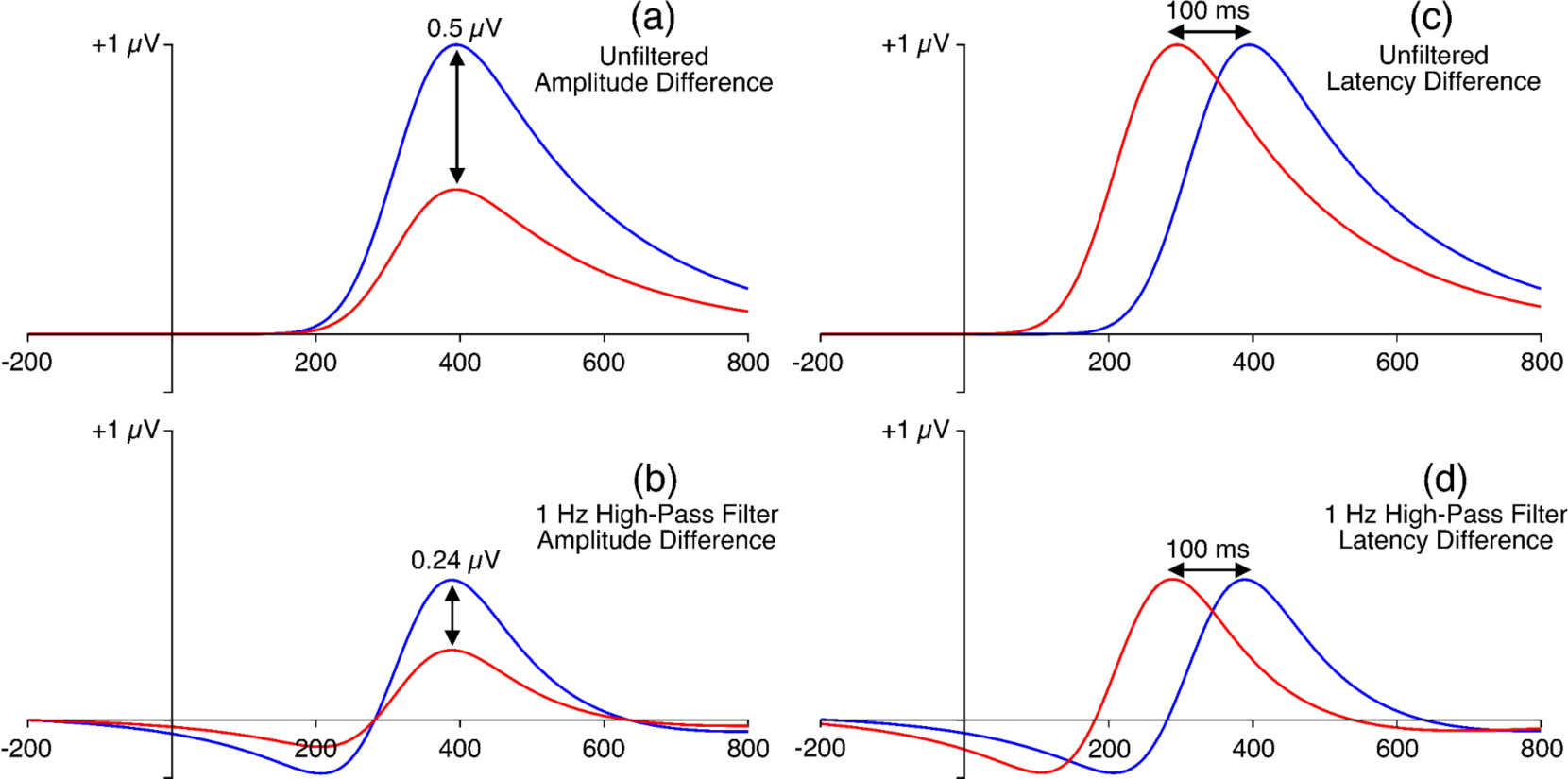
Demonstration of how filters reduce amplitude differences but not latency differences. (a) Simulated P3 wave in two conditions, with a difference in peak amplitude of 0.5 µV between the conditions. (b) Same waveforms as in (a) after the application of a high-pass filter with a 1 Hz half-amplitude cutoff and a slope of 12 dB/octave. The filtering caused a reduction in the amplitude difference to 0.24 µV. (c) Simulated P3 wave in two conditions, with a difference in peak latency of 100 ms between the conditions. (d) Same waveforms as in (c) after the application of a high-pass filter with a 1 Hz half-amplitude cutoff and a slope of 12 dB/octave. The difference in latency between the two conditions is still 100 ms. The waveforms were created using ex-Gaussian functions with a standard deviation of 58, a tau of 2000ms, an amplitude of 0.5 µV or 1µV, and a Gaussian mean of 320ms or 220ms.

Filters do not typically have large effects on latency scores, and any observed effects may consist of an increase or a decrease depending on the shape of the waveform. Moreover, if a shift in latency does occur, the latency scores will typically be shifted equivalently across conditions. This is illustrated in Figures 10c and 10d, which show a simulation of two conditions in which the P3 latency differs. In the unfiltered data (Figure 10c), the peak latency differed by 100 ms between the two conditions. When a high-pass filter with a half-amplitude cutoff of 1 Hz was applied (Figure 10d), the peak latency was shifted leftward by 6.4 ms in both conditions, and the difference in peak latency between the two conditions remained 100 ms. Because filtering does not consistently reduce the difference in latency between groups or conditions, it is not typically necessary to consider the effects of filtering on the signal relative to the noise (the SNR) when selecting an optimal filter for latency scores. Indeed, the SNR might even be misleading, because a filter might lead to smaller (earlier) latency scores even if it does not decrease the ability to detect differences between groups or conditions.

Instead, one can simply determine which filters yield the lowest noise (the smallest SME value), along with a consideration of waveform distortion. Thus, the flowchart shown in Figure 2 would be altered slightly for latency scores, using RMS(SME) rather than SNR_SME_ to assess data quality and choosing the filter that produces the lowest RMS(SME) while also producing an APP of less than 5%.

### 7.1. Example: Selecting optimal filter parameters for P3 peak latency

In this subsection, we will illustrate our process of selecting filter parameters for latency scores using the same dataset used for the P3 mean amplitude example. We will focus on P3 latency, scored as the peak latency between 300 and 600 ms in the target-minus-standard difference wave at the Pz electrode site We examined the same set of candidate filters as in our analyses of P3 mean amplitude.

In the data quality quantification portion of our procedure (Figure 2, left), we again looped through the candidate filters, computing a target-minus-standard difference wave for each participant for a given filter. We used bootstrapping to obtain the SME for P3 peak latency for that filter (see Section S6 of the supplementary materials for details). We then computed the RMS of the single-participant SME values as our overall estimate of the noise level for that filter. The resulting RMS(SME) values are shown in Figure 11 (see supplementary Figure S1 for the grand average waveforms).

**Figure 11.**
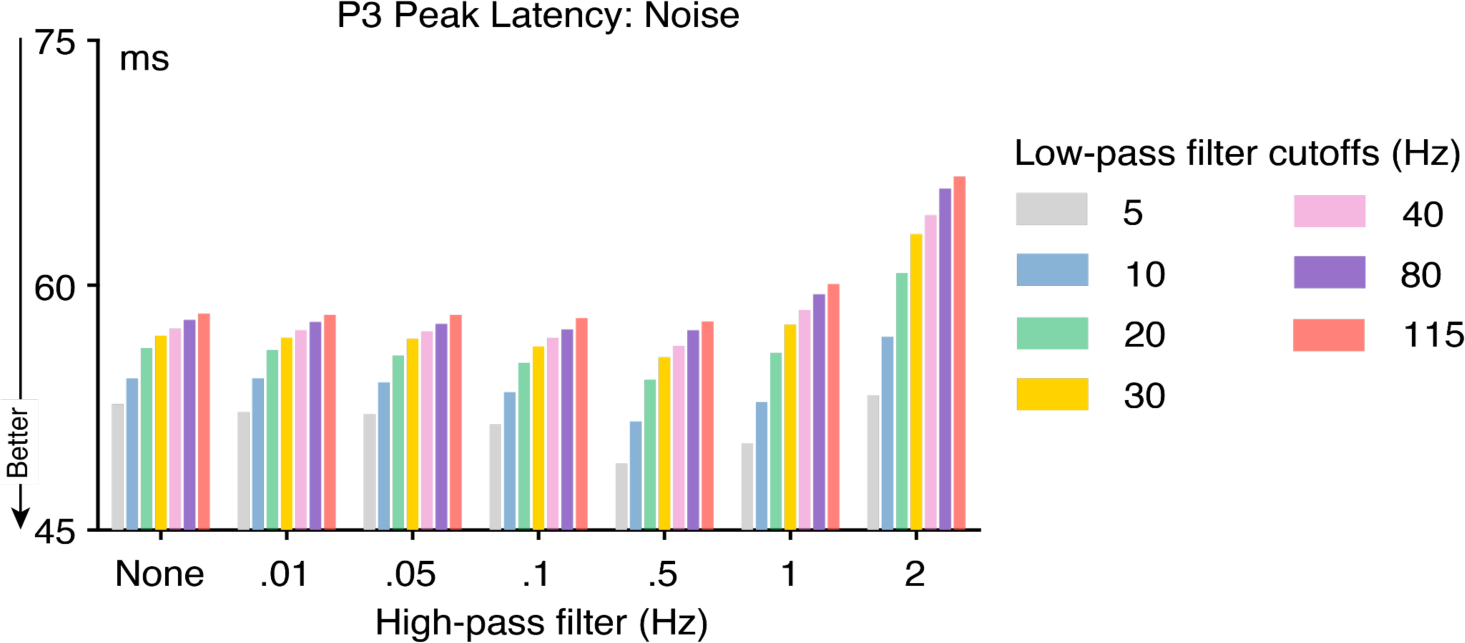
Noise defined by root mean square of the standardized measurement error values for P3 peak latency from the ERP CORE across a wide range of filter combinations. The continuous EEG was filtered prior to averaging with every combination of seven high-pass filter cutoffs (0, 0.01, 0.05, 0.1, 0.5, 1, and 2 Hz) and seven high-pass filter cutoffs (5, 10, 20, 30, 40, 80, and 115 Hz). All filters used here were noncausal Butterworth filters with a slope of 12 dB/octave, and cutoff frequencies indicate the half-amplitude point.

The high-pass cutoff frequency had relatively little impact on the RMS(SME) until the cutoff reached extreme values (1 Hz or greater). By contrast, the low-pass cutoff frequency had a large impact, with progressively smaller (better) RMS(SME) values as the cutoff frequency decreased. This is the opposite of the pattern that was observed for mean amplitude scores, where the high-pass cutoff had a large effect on the RMS(SME) and the low-pass cutoff had little or no effect (see Figure 9b). These opposite patterns reflect the fact that mean amplitude scores are strongly impacted by low-frequency noise but not by high-frequency noise, whereas peak latency scores are strongly impacted by high-frequency noise but not by low-frequency noise. This further reinforces the general point that the optimal filter settings depend on how the data will ultimately be scored.

The next step was to assess waveform distortion using the artifactual peak percentage (APP). This is done in exactly the same way for latency measures as for amplitude measures, so we used the APP values that we obtained for our analyses of P3 mean amplitude in Section 6. We then combined this information with the RMS(SME) information to select the optimal filter.

The best RMS(SME) value was produced by the combination of a 0.5 Hz high-pass filter and a 5 Hz low-pass filter, but all the filter combinations including a 0.5 Hz high-pass filter exceeded the 5% APP threshold. Of the filters with an APP less than 5%, the best RMS(SME) was produced by the combination of a 0.1 Hz high-pass filter and a 5 Hz low-pass filter.

We also examined a denser sampling of high-pass cutoffs (see supplementary Figure S3 for the RMS(SME) and APP values). Of the filters with an APP of less than 5%, the lowest RMS(SME) value was obtained for the combination of a 0.2 Hz high-pass filter and a 5 Hz low-pass filter. This is therefore the optimal filter when P3 peak latency is scored from studies like the ERP CORE visual oddball experiment. Note that this is a much lower low-pass cutoff frequency than the 30 Hz cutoff that we previously recommended for general use in cognitive and affective research (Luck, 2014). This shows the value of using a formal procedure to determine the optimal filtering parameters.

A 5 Hz low-pass filter produced a modest but clearly noticeable leftward shift in the onset latency of the simulated P3 (see the upper left panel in Figure 7). If this latency distortion might impact the scientific conclusions of a given study, a higher cutoff frequency could be chosen for the low-pass filter. For example, increasing the low-pass cutoff from 5 Hz to 10 Hz reduces the leftward shift in the onset latency while only increasing the RMS(SME) by less than 5% (from 50.1 to 52.4; see supplementary Figure S3).

## 8. More complex scenarios

Our approach to filter selection was based on a simple scenario, in which the researcher wishes to determine the optimal filter for a single score (e.g., P3 peak latency) and the component of interest can be well isolated by means of a difference wave (e.g., target-minus-standard). This section considers more complex scenarios.

### 8.1. Analyzing multiple scores in a given study

One likely scenario is that a researcher is interested in more than one score in a given study (e.g., both peak amplitude and peak latency for the P3 component, or mean amplitude for both the N170 and P3 components). If different filters are optimal for the different scores, what filter settings should be used? This will depend on the scientific goals of the experiment. In most cases, the most conservative approach would be to use the same filter settings for all the scores in a given study. In this case, it would be typically appropriate to choose a filter that strikes a reasonable compromise between the data quality for the different scores while ensuring that the filter does not exceed the threshold for waveform distortion for any of the components.

However, there may be good reasons to use different filters for different components in a single study when the components have very different frequency properties. For example, Woldorff et al. (1987) examined the effects of selective attention on the very rapid auditory brainstem responses (ABRs, <10 ms), the auditory midlatency responses (MLRs, 20-50 ms), and the auditory long-latency responses (LLRs, >80 ms). These components have very different frequency properties, so the researchers one bandpass for the ABRs (30-2000 Hz) and a very different bandpass for the MLRs and LLRs (0.01-100 Hz). The goal of that study was to determine whether attention impacted each of these responses, and using the best filter settings for each component was a reasonable way of maximizing the ability to detect attention effects for the different components.

In addition, different filter settings are often optimal for different scoring methods. For example, the companion paper (Zhang et al., 2023) found little or no impact of the low-pass filter cutoff on data quality for P3 mean amplitude scores but found that a 5 Hz low-pass cutoff improved the data quality for P3 peak amplitude. Onset latencies are particularly prone to distortion from noise. Thus, researchers sometimes use lower low-pass cutoffs for latency measures than for mean amplitude in the same study. For example, Kang et al. (2019) used a low-pass cutoff at 30 Hz for their amplitude analyses but used a cutoff at 8 Hz for their analyses of the onset latency of the lateralized readiness potential. In general, researchers must think carefully about the consequences of using identical or different filters for different scores in the same dataset, and they should explicitly justify their choices.

### 8.2. Temporally adjacent components

ERP waveforms typically contain a progression from relatively small and narrow components (e.g., P1) to relatively large and broad components (e.g., P3). When multiple components are impacted by a given experimental manipulation, the difference wave may contain a smaller, narrower component followed by a larger, broader component. For example, a target-minus-standard difference wave in an oddball experiment often contains a relatively small and narrow N2 followed by a large and broad P3. When this happens, a filter that is appropriate for one of the components may produce significant waveform distortion for the other component, and artifactual effects may occur if the wrong filter is applied.

This is illustrated in Figure 12, which illustrates the effects of filtering a simulated waveform that contains both an MMN component and a P3 component (based on the waveform parameters for the simulated MMN and P3 waveforms in the companion paper). Figure 12a shows the waveform for the simulated MMN, along with the waveform after application of a 0.5 Hz high-pass filter (which was the optimal cutoff for the MMN and produced minimal waveform distortion). Figure 12b shows the simulated P3, both unfiltered and filtered at 0.5 Hz. Although this filter was optimal for the MMN, it produced a substantial distortion of the P3, including a negative-going deflection during the time period of the MMN. When both the MMN and P3 were summed together into a single waveform (Figure 12c), the filtered waveform appeared to contain a larger MMN because of the negative-going filter artifact for the P3. Figures 12d and 12e show how this can lead to an incorrect interpretation. Figure 12d shows the unfiltered waveforms for two conditions, one in which the P3 is 50% larger than in the other. Figure 12e shows the result of filtering these waveforms. The artifactual negativity produced by the filter during the MMN time period is now larger for the condition with the larger P3 wave, making it appear as if the both the MMN and the P3 were larger in this condition (see Acunzo et al., 2012; Tanner et al., 2015, for similar artifactual effects).

**Figure 12.**
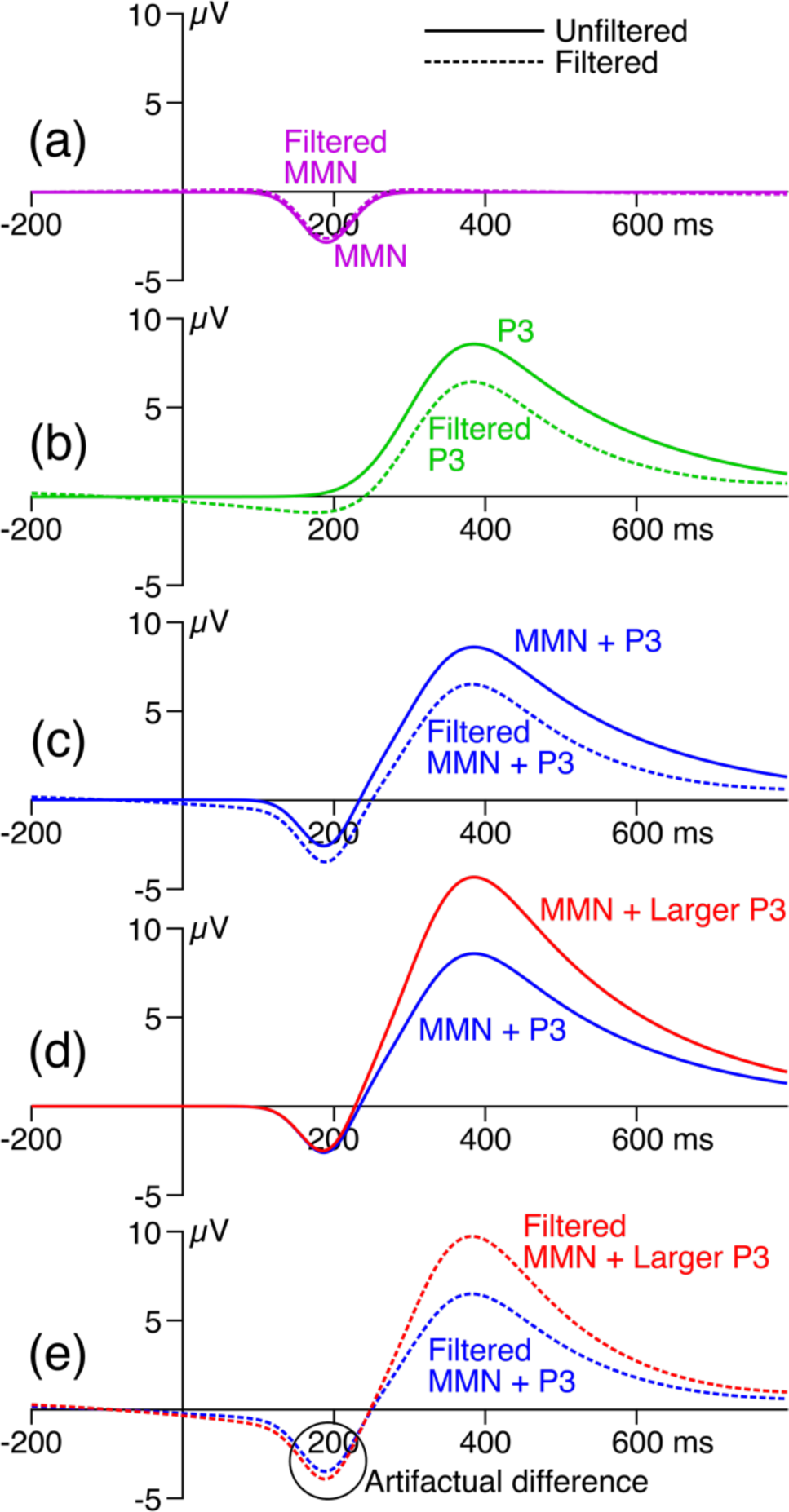
Effects of filtering a simulated waveform that contains both a mismatch negativity (MMN) and a P3. (a) Waveform for the simulated MMN, along with the waveform after application of a 0.5 Hz high-pass filter (12 dB/ octave). (b) Waveform for the simulated P3, along with the waveform after application of a 0.5 Hz high-pass filter (12 dB/ octave). (c) Summation of the MMN and P3 waveforms into a single waveform. (d) Unfiltered waveforms for two conditions, one in which the P3 is 50% larger than in the other. (e) Filtered waveforms for the two conditions (0.5 Hz high-pass filter, 12 dB/octave).

This problem can be minimized by choosing a filter that produces minimal waveform distortion (e.g., APP < 5%) for each of the individual components. To be even more careful (or when the waveshapes of the underlying components are not known), researchers could create an artificial waveform that reflects the complex shape of the observed waveform and examine how this waveform is distorted by the filter (as in Figure 12c).

### 8.3. Temporally overlapping components

An even more challenging problem arises when multiple components are strongly active in the same time period. In a typical ERP waveform, the voltage at a given electrode site at a given latency reflects a weighted sum of multiple different underlying components (Coles & Rugg, 1995; Donchin & Heffley, 1978; Luck, 2014). For example, the voltage produced by a target stimulus at the Pz electrode site at 400 ms may reflect the combined impact of a dozen different ERP components, not just the P3 component. A given filter may change the data quality differently for these different components, and a filter may also produce different waveform distortion patterns for the different components as a result of their different waveshapes. This makes it difficult to find a filter that is optimal for all the components that are active in the ERP waveform.

In the present work, we have used a common solution to this problem, namely using difference waves to subtract away most of the components, ideally leaving only a single component in the waveform. Difference waves work reasonably well in many cases, but they often fail to isolate a single component. For example, a target-minus-nontarget difference wave often includes an N2 component as well as a P3 component (Folstein & Van Petten, 2008), and the P3 portion may include multiple different components (Polich, 2012). In addition, difference waves may not be appropriate for assessing some kinds of effects, such as the error-related negativity that is present on non-error trials (Gehring et al., 2012). Researchers may choose to use other approaches to isolating components in these situations, such as source reconstruction (Michel & Brunet, 2019) or spatiotemporal principal component analysis (Spencer et al., 2001). It would be straightforward to insert these methods into our procedure for determining the optimal filter settings.

## 9. Limitations and future directions

The present approach to filter selection has several strengths, including the use of objective and quantifiable properties of filters with respect to specific ERP effects and the scoring methods used to quantify them. However, subjective decisions are involved in selecting the shape of the artificial waveforms that are used to assess waveform distortion. In addition, the 5% artifactual peak percentage criterion, although reasonable, is somewhat arbitrary. Nonetheless, the present approach makes it possible to quantitatively assess both the benefits and costs of filtering.

Another important limitation of the present approach is that it is designed only for studies in which the scores are obtained from averaged ERP waveforms (because the SME is a measure of data quality for such scores). It is not designed for single-trial analyses, which can be quite valuable and are becoming increasingly common (Bürki et al., 2018; Heise et al., 2022; Volpert-Esmond et al., 2018; Winsler et al., 2018). However, there is good reason to believe that the present approach will work well for single-trial analyses in which mean amplitude is scored from single-trial EEG epochs and these single-trial scores are then entered into the statistical analyses. This scoring method is a linear operation, as is averaging across trials, and the filters typically used in ERP research also involve a linear or approximately linear operation. The order of operations does not matter for linear operations (Luck, 2014), so the effects of filtering should be the same whether the mean amplitude is measured before or after averaging. Thus, we conjecture that our filter selection approach will also be well suited for single-trial analyses using mean amplitude scores. However, additional research would be required to verify this conjecture.

The present approach is also not designed for mass univariate analyses, in which statistical comparisons between groups or conditions are made at a large number of individual time points and/or electrode sites, accompanied by an appropriate correction for multiple comparisons (Frossard & Renaud, 2022; Groppe et al., 2011; Maris & Oostenveld, 2007). As discussed in Section 3.2, the SNR for an individual time point can be estimated by simply dividing the voltage at that time point by the standard error of the mean at that time point. Rather than using SNR_SME_, this traditional SNR value could be used in selecting filter parameters. However, it is not obvious how one would combine standard errors across time points and/or electrode sites or how these standard error values would interact with the procedure for correcting for multiple comparisons. This is another avenue for future research.

## Acknowledgments

We thank Dr. Lindsay Bowman for excellent advice about how to make the filtering approach described in this manuscript useful across a broad range of research areas, recording methods, and participant populations.

## Supplementary Materials

### S1. Table S1

**Table S1:**
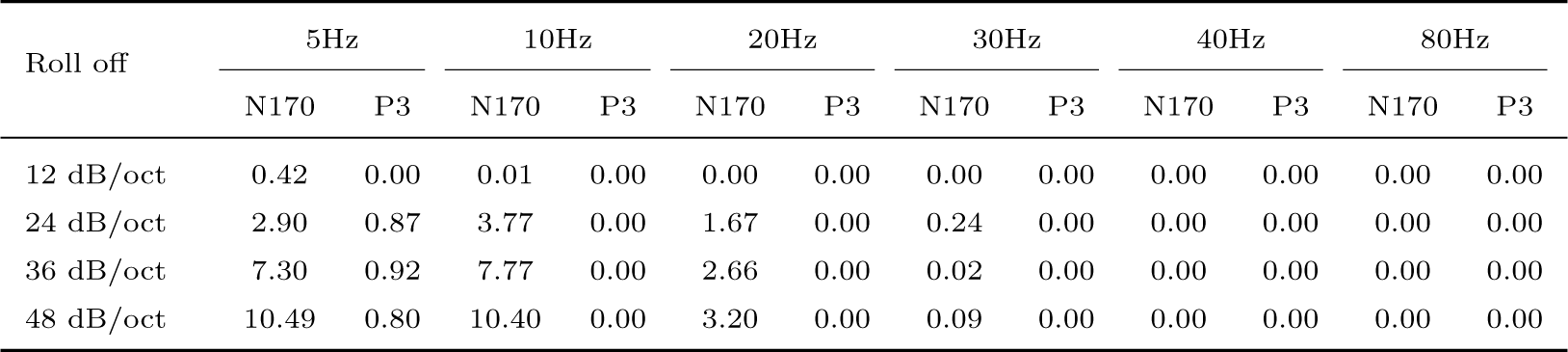
Artifactual peak percentage values (in %) for low-pass filters with different roll-offs (12, 24, 36, and 48 dB/octave) for simulated N170 and P3 waves.

| Roll off | 5Hz |  | 10Hz |  | 20Hz |  | 30Hz |  | 40Hz |  | 80Hz |  |
| --- | --- | --- | --- | --- | --- | --- | --- | --- | --- | --- | --- | --- |
|  | N170 | P3 | N170 | P3 | N170 | P3 | N170 | P3 | N170 | P3 | N170 | P3 |
| 12 dB/oct | 0.42 | 0.00 | 0.01 | 0.00 | 0.00 | 0.00 | 0.00 | 0.00 | 0.00 | 0.00 | 0.00 | 0.00 |
| 24 dB/oct | 2.90 | 0.87 | 3.77 | 0.00 | 1.67 | 0.00 | 0.24 | 0.00 | 0.00 | 0.00 | 0.00 | 0.00 |
| 36 dB/oct | 7.30 | 0.92 | 7.77 | 0.00 | 2.66 | 0.00 | 0.02 | 0.00 | 0.00 | 0.00 | 0.00 | 0.00 |
| 48 dB/oct | 10.49 | 0.80 | 10.40 | 0.00 | 3.20 | 0.00 | 0.09 | 0.00 | 0.00 | 0.00 | 0.00 | 0.00 |

### S2. Figure S1

**Figure S1:**
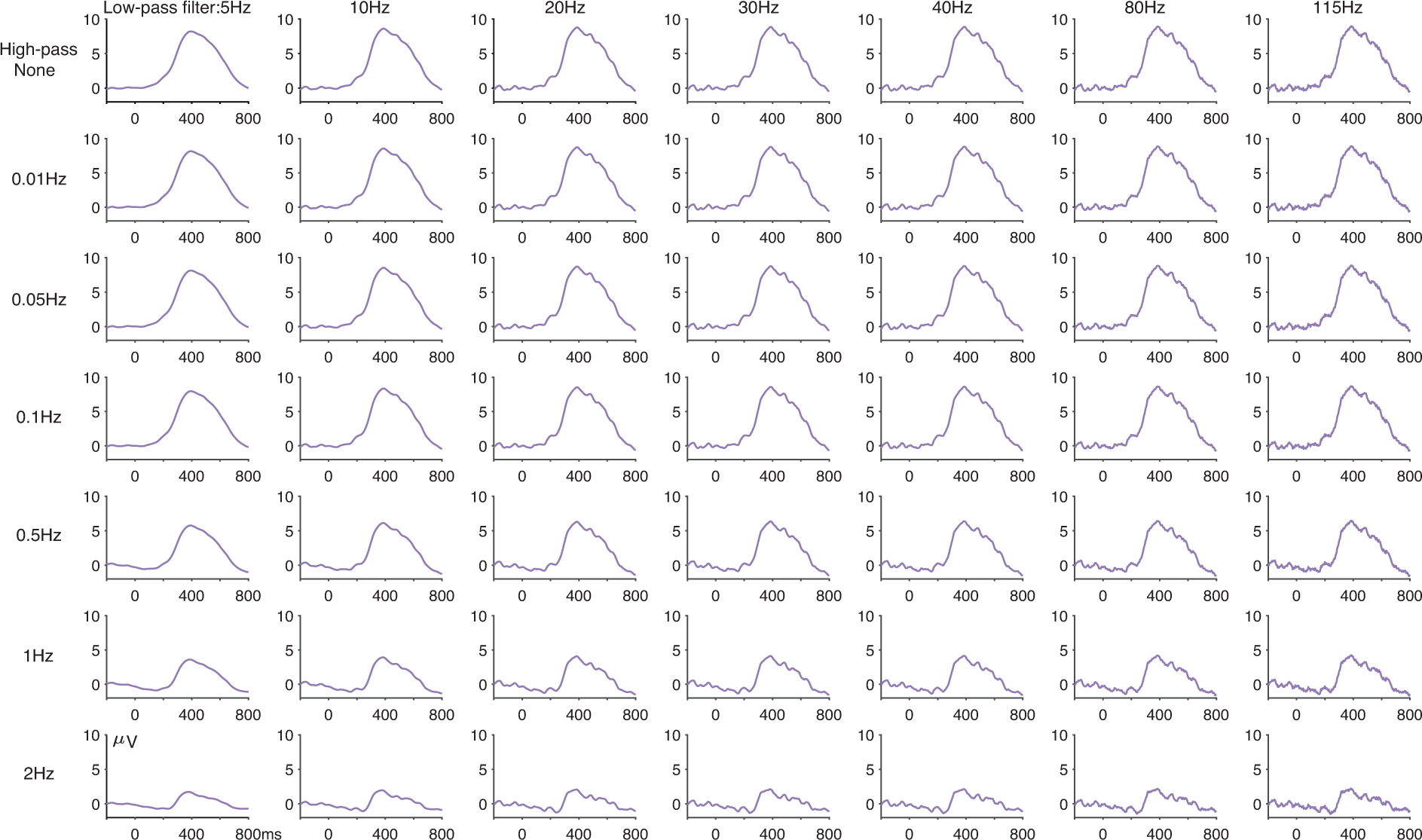
Grand average target-minus-standard difference waves from the ERP CORE oddball experiment at the Pz electrode sit for each combination of seven high-pass filter cutoffs (0, 0.01, 0.05, 0.1, 0.5, 1, and 2 Hz) and seven high-pass filter cutoffs (5, 10, 20, 30, 40, 80, and 115 Hz). All filters used here were noncausal Butterworth filters with a slope of 12 dB/octave, and cutoff frequencies indicate the half-amplitude point.

### S3. Figure S2 and the effects of filtering on scalp topography

**Figure S2:**
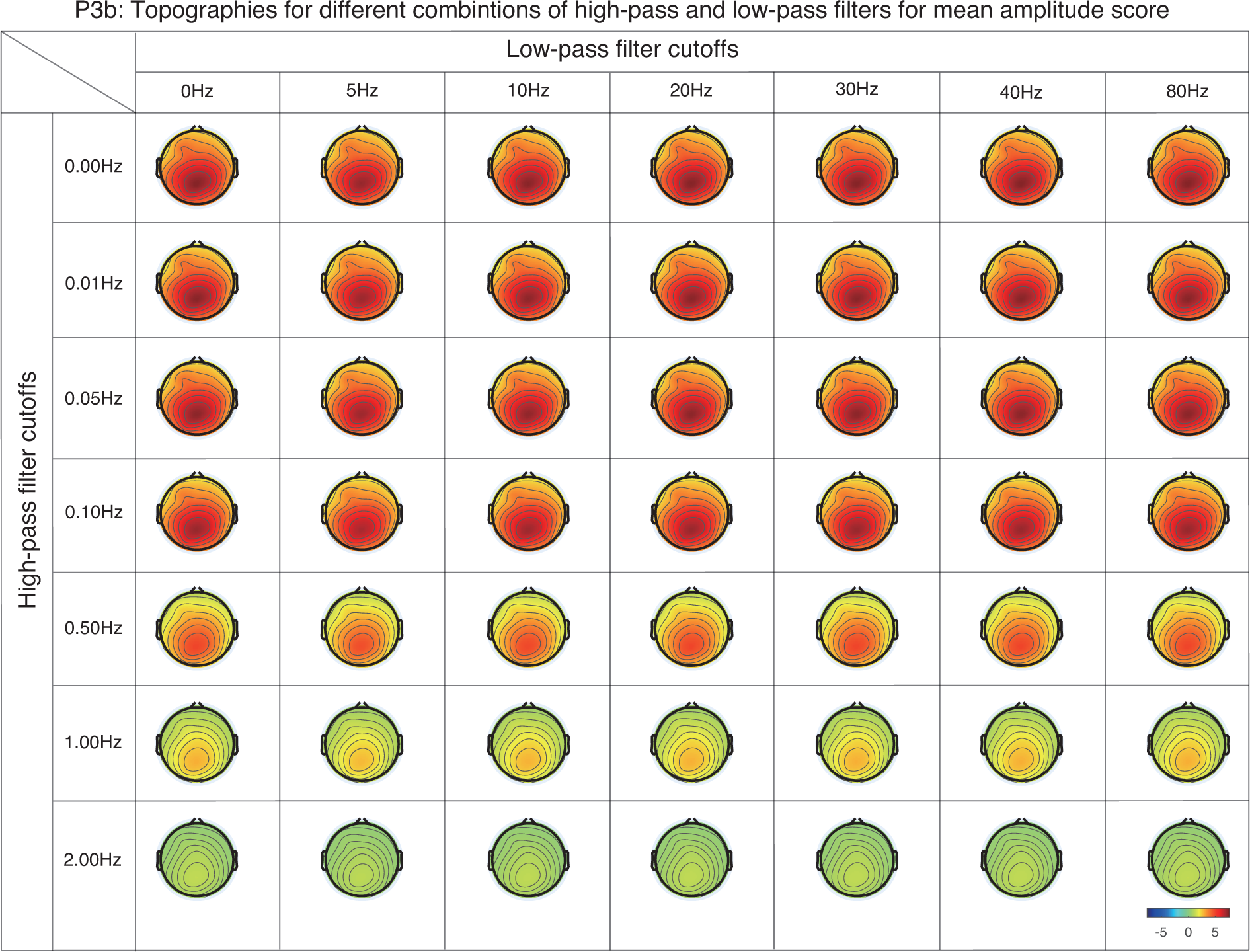
Filtering and scalp topography

The P3 analyses presented in Section 6 of the main manuscript were obtained from the Pz electrode site, which was the a priori measurement site for the ERP CORE P3 experiment. In many studies, multiple electrode sites are averaged together into a cluster prior to measuring the amplitudes or latencies, which can improve the data quality (Zhang & Kappenman, 2023). The present approach can be applied to values obtained from these clusters (but note that the clusters must be created prior to averaging to apply ERPLAB’s automated SME estimation routine to the clusters). In other studies, however, amplitude or latency values are obtained separately from each of many sites (e.g., for mass univariate analyses). It is therefore natural to ask how filtering will impact the scalp topography of the ERP signal.

If a single component has been well isolated using a difference wave (or some other approach), then the unfiltered waveform at each electrode site will be a scaled copy of the waveform at the maximal site (because ERPs propagate multiplicatively; McCarthy & Wood (1985)), plus or minus any site-specific noise. Filtering the data will change the waveform at the maximal site, and the filtered waveforms at the other sites will be scaled copies of this new waveform, plus or minus any site-specific noise. As a result, the scalp topography will not be changed by filtering under these conditions. This is illustrated in this figure, which shows that the topography of the P3 difference wave was approximately the same for the different combinations of low-pass and high-pass filter cutoffs (except for differences in amplitude). However, this equivalence may not hold when multiple components with different topographies are strongly and simultaneously active in the difference wave, in which case simulations may be necessary to predict the results.

It is possible that the frequency content of the noise would vary systematically across electrode sites. If so, then the best *SNR_SME_* might arise from different filter settings at different electrode sites. Although we have not formally tested this situation, we anticipate that these differences would be modest under typical EEG recording conditions (e.g., the best high-pass cutoff being 0.2 Hz at some sites and 0.3 Hz at other sites, with modest differences in *SNR_SME_* between these cutoffs). If this situation arises, a simple solution would be to “split the difference” and select a cutoff at the mean of the optimal values for the different sites (e.g., a high-pass cutoff of 0.25 Hz). However, additional research would be necessary to rigorously assess these issues.

### S4. Figure S3

**Figure S3:**
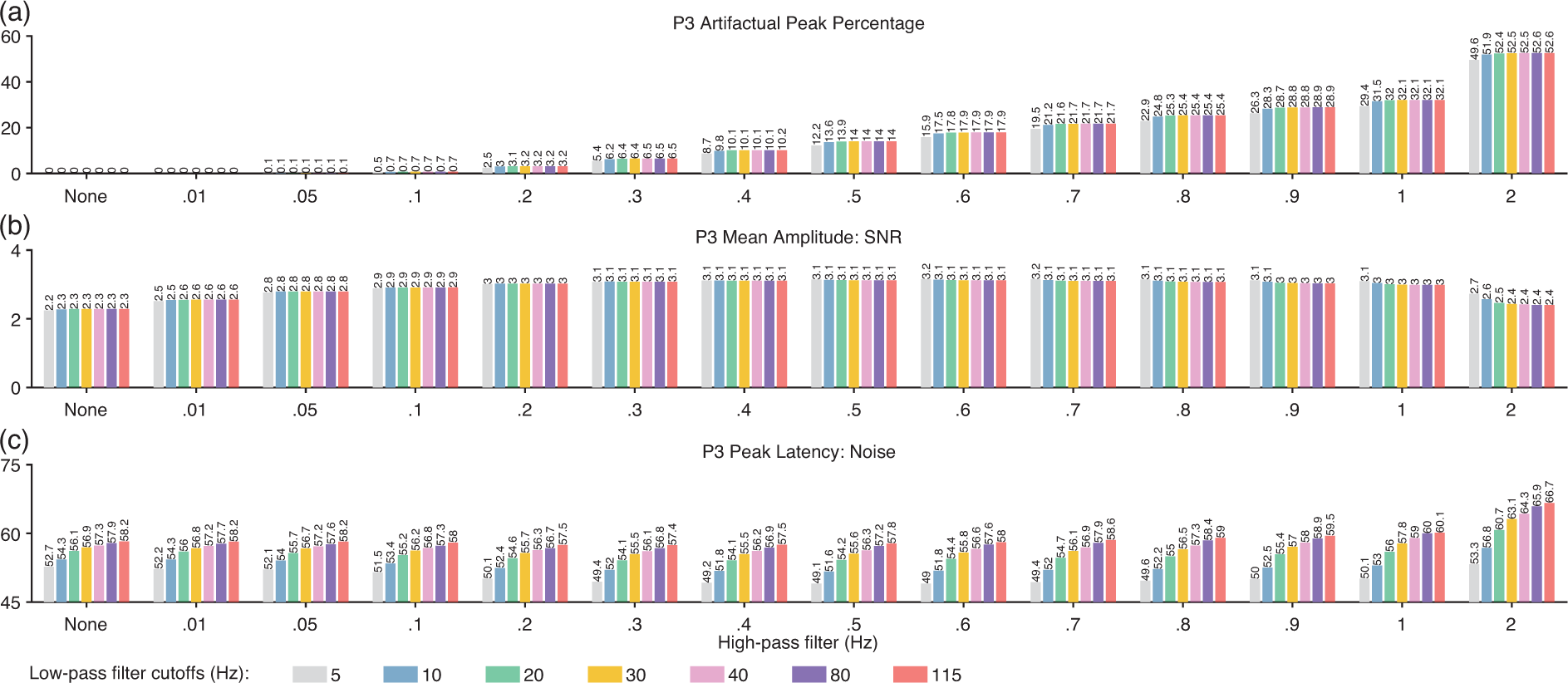
Metrics of filtering effectiveness (a) and artifacts for P3 mean amplitude (b) and peak latency (c) for each combination of high-pass filter cutoffs (0, 0.1, 0.2, 0.3, 0.4, 0.5, 0.6, 0.7, 0.8, 0.9,1, and 2Hz) and low-pass filter cutoffs (5, 10, 20, 30, 40, 80, and 115 Hz). (a) Artifactual peak percentage. (b) Signal-to-noise ratio for P3 mean amplitude. (c) Noise level (standardized measurement error) for P3 peak latency. All used filters here were noncausal Butterworth filters with a slope of 12 dB/octave, and cutoff frequencies indicate the half-amplitude point.

### S5. Essence of the standardized measurement error (SME)

The SME quantifies the standard error of measurement for a given amplitude or latency score. Roughly speaking, it indicates our confidence (or lack of confidence) that the score obtained for a given participant is representative of that participant’s true score. More precisely, the SME — like any other standard error — quantifies the degree to which a given score would be expected to vary if we repeated the experiment a very large or infinite number of times in a given participant (assuming no fatigue or learning) and measured the score for each repetition of the experiment.

In other words, the standard error of a score for a given participant reflects what would happen if, in theory, we repeated an experiment many times with that participant and obtained the score of interest for each repetition of the experiment (assuming no learning, fatigue, etc.). For example, we could run an oddball task an infinite number of times for a given participant, obtain the target-minus-standard difference wave for each repetition, and get the peak latency from this difference wave for each repetition. This would yield an infinite number of peak latency scores, and the true score for that participant would be the mean of this infinite set of scores. The standard error of the peak latency for that participant would be the standard deviation of that infinite set of peak latency scores. For more information about standard errors and other similar measures (see Cronbach & Shavelson (2004); Hedge et al. (2018); Williams et al. (1995); Williams & Zimmerman (1989)).

We cannot, in practice, repeat an experiment an infinite number of times for a given participant. Even if we could repeat an experiment a large number of times, the scores would change systematically over repetitions as a result of factors such as learning and fatigue. Fortunately, we can estimate the standard error from the data obtained in a single experiment for a given participant. When we scoring the mean amplitude (i.e., the mean voltage within a specific time range), we can use the equation for the standard error of the mean (which is a special case of the standard error of measurement). That is, we get a mean amplitude score on each trial, take the standard deviation of these single-trial scores, and divide by the square root of the number of trials:

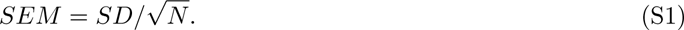

This equation is not valid for other scores (e.g., peak amplitude, peak latency), but we can use bootstrapping to estimate the standard errors for these scores (as described in the next section). The SME is then equal to these standard errors. Both Equation S1 and bootstrapping make the assumption that the data are completely independent on the different trials of an experiment. This assumption is almost certainly incorrect because the trials are obtained sequentially. There may be autocorrelations (e.g., a larger-than-normal value on trial N when there is a larger-than-normal value on trial N-1) and there may be gradual trends (e.g., progressively smaller amplitudes over the course of the session). These issues are discussed in detail in Section S8 of the supplementary materials for Luck et al. (2021), and potential solutions are provided. For the present purposes, these issues are likely to be negligible. The autocorrelations are likely to be small once the data have been baseline-corrected. Gradual trends are more likely, but they will tend to be small in short experiments (such as the ERP CORE experiments, each of which lasts approximately 10 minutes). Moreover, the present goal is to compare relative differences in data quality across different filter settings, not the absolute data quality. Thus, violations of the assumption of independence are unlikely to have a major effect when the SME is used in the context of assessing the data quality resulting from different filters. In experiments with large autocorrelations (e.g., experiments without baseline correction) or experiments with large gradual drifts in the amplitudes or latencies (e.g., very long experiments that might produce substantial learning or fatigue), the solutions described in Section S8 of the supplementary materials for Luck et al. (2021) can be used. It should be noted that there are two different classes of measurement error, random errors and systematic errors (see Brandmaier et al. (2018) for a detailed discussion). These are quantified as the precision of a measure (the extent to which the values are similar across repeated measurements) and the bias of a measure (the extent to which the average value is consistently less than or greater than the true value). The SME reflects the precision of an ERP score, not the bias. Filters also create biases by reducing amplitudes and creating artifactual peaks. We assess these biases separately from the SME.

### S6. Using bootstrapping to estimate the SME

Bootstrapping is a widely used approach for estimating the standard error of measurement (Boos, 2003; Efron & Tibshirani, 1994). A general introduction to bootstrapping in the context of ERPs is provided in Section S4 of the supplementary materials for Luck et al. (2021). Here we provide a more condensed description of our approach and a brief discussion of the limitations of this approach.

As described in the preceding section, the standard error of a score for a given participant reflects the variability (standard deviation) in the score that would be expected across a large or infinite number of replications of the experiment in that participant. We typically have the data from only one run of the experiment for a given participant, but we can use a standard statistical “trick” to simulate running a very large number of experiments for this participant. The trick is to randomly sample from the trials that were actually obtained to simulate a new experiment. For example, imagine that we have 40 target trials and 160 standard trials in a P3 experiment for a given participant. To simulate a new experiment for this participant, we randomly select 40 of the target trials and 160 of the standard trials from the actual set of trials, sampling with replacement, and then we make averaged ERPs for the target and standard from these randomly sampled trials. We then make a target-minus-standard difference wave and get the peak P3 latency score from this difference wave. We then repeat this a total of 1000 times, which gives us 1000 peak latency scores for this participant. The SME is then the standard deviation of these 1000 scores.

Note that it is necessary to sample with replacement from the set of available trials. Otherwise the averaged ERP waveforms will be the same for each repetition. In addition, it is necessary to have the same number of sampled trials for each simulated experiment be the same as the number of trials that are being averaged together in each condition in the actual experiment (because the noise level in the averaged ERP depends on the number of trials being averaged together). This is based on the number of trials being included in the averaged ERP waveforms used in the actual experiment. For example, if participants received 40 target trials, but a given participant had only 31 trials after the rejection of trials with artifacts and behavioral errors, the bootstrapping would be accomplished by randomly sampling 31 of these 31 trials (with replacement). As a result, the noise level of the averaged ERPs in the simulated experiments will reflect the number of trials included in the main analyses for a given participant. In general, the analyses for the simulated experiments must be identical to the main analyses except that the trials for the simulated experiments are sampled (with replacement) from the set of trials used in the main analyses.

### S7. Quantifying latency distortion

For cases in which temporal smearing should be considered in selecting filter settings, we recommend quantifying the amount of smearing with a latency distortion percentage that is analogous to the artifactual peak percentage. This would involve quantifying the width of the filtered and unfiltered artificial waveforms as the full width at half maximum (FWHM). The FWHM is computed by finding the time points on the two sides of the peak at which the amplitude is 50% of the peak amplitude and calculating the difference in latency between these points. The latency distortion percentage is then calculated as the absolute value of the FWHM for the filtered waveform divided by the absolute value of the FWHM for the filtered waveform. The threshold for an acceptable latency distortion percentage will depend on the specific research question.

### S8. SME for differences between conditions

When the amplitude of an ERP component is scored as the mean voltage within a time range, the SME for a single experimental condition can be calculated analytically using Equation 1. This equation involves obtaining the mean voltage score from each single-trial epoch, computing the standard deviation (SD) of these scores, and dividing by the square root of the number of trials. This approach is fast and straightforward, and it is directly implemented in ERPLAB Toolbox. However, many ERP experiments use a difference wave to isolate an ERP component (e.g., target minus standard for the P3, related minus unrelated for the N400). Indeed, researchers will often score an amplitude or latency from a difference wave and enter these scores into the statistical analyses. In such cases, it is valuable to quantify the data quality for the score obtained from the difference wave. It is possible to do this via bootstrapping, but bootstrapped SME values take much longer to compute and currently require custom scripting. An alternative is to obtain the analytic SME for each of the individual conditions using Equation S1 and use Equation S2 to estimate the SME for the difference score using Equation S2:

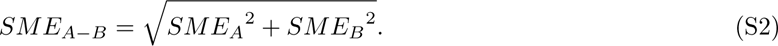

In this equation, *SME_A-B_* is the SME of the difference between conditions *SME_A_* and *SME_B_*, and *SME_A_* and *SME_B_* are the SMEs of the two individual conditions (obtained using Equation S1). Here, we explain why Equation S2 is valid in this situation.

Equation S2 is based on the equation for the variance between the sum or difference between two independent random variables A and B:

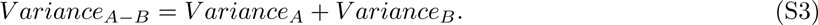

SME values are a type of standard deviation, so we can rewrite this equation in terms of SDs:

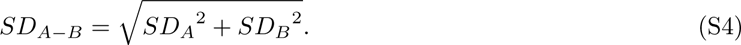

This is essentially the same as Equation S2. However, these equations assume that A and B are independent of each other, and this may seem like an invalid assumption. For example, imagine that we obtained the mean amplitude of the P3 wave for targets and standards in an oddball experiment. If we obtained these scores for each participant, they would not be independent variables. That is, participants with a larger-than-average score for the standards might also be expected to have a larger-than-average score for the targets. In other words, the two variables are correlated. The equation for estimating the variance of the sum or difference of two correlated variables is:

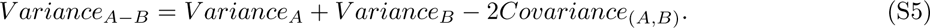

In this situation, we have pairs of A and B values, with one pair of values each participant. However, this is not a good analogy for the SME of the difference between conditions, because the SME is computed separately for each individual participant. For a given participant, we do not have pairs of A and B values. Instead, we have a sample of A values (one for each trial in condition A) and a sample of B values (one for each trial in condition B). There is no pairing of a given A value with an individual B value. So there is no covariance between A and B.

In other words, we are examining the variability across trials within a single participant, which can be thought of as samples from an infinite population of trials for that participant. The trials do occur in a given order, and the score on one trial may provide some information about the score on the next trial (i.e., an autocorrelation). This would be like testing participants in two groups, A and B, in a random order. For example, we might have someone in group A on one day, someone in group B on another day, another person in group B the next day, etc. There might be some autocorrelation across days, but this creates a covariance across days rather than a covariance across groups. Consequently, we would simply apply Equation S1 to estimate the standard error of the mean for a given group. And if we wanted to estimate the standard error of the difference in means between the two groups, we would treat the means for groups A and B as independent variables and apply Equation S2. Thus, estimating the SME for a difference score is analogous to estimating the standard error of the difference between means in two separate groups of subjects, and Equation S2 is appropriate.

Moreover, we have verified that the SME values obtained using Equation S2 are nearly identical to the values obtained using bootstrapping. Thus, it appears to provide an excellent approximation in practice. Researchers who are uncomfortable with the assumption of independence can instead use bootstrapping to more directly assess the SME for scores obtained from difference waves. Note that Section of the supplementary materials for Luck et al. (2021) contains a more extensive discussion of the issue of autocorrelation across trials.

There are two important exceptions to Equation S2. First, it applies only when the score is the mean voltage across a time window (because this is the only situation in which Equation S1 can be used to obtain the SME from each condition). Second, it applies only when the difference wave is between waveforms based on separate trials, not when it is a difference between two electrode sites (e.g., contralateral minus ipsilateral). When A and B are scores obtained from different electrode sites on the same trial, they are paired, and the covariance between them would need to be taken into account when estimating the SME of the difference between them.

## Footnotes

1 The general idea of choosing filters that maximize the data quality and avoid waveform distortion has been explored in the context of auditory sensory responses, but with quantification approaches that reflect the specific issues involved in that domain (e.g., identifying whether a sensory response was present). See Picton (2000) for a review.

2 Many different filtering algorithms are available, but for simplicity we focus on noncausal Butterworth filters (Hamming, 1998). This class of filters was chosen because it is efficient, flexible, well-behaved, and widely used for EEG and ERP signals. However, our general approach is independent of the filtering algorithm.

3 An ex-Gaussian function is a Gaussian function convolved with an exponential function to create a skewed waveform. This function is often used to model reaction time distributions (e.g., Karalunas et al., 2014; Schmiedek et al., 2007), which are typically right-skewed. Long-latency ERP waves are also typically right-skewed, often as a result of the same factors that cause reaction time variability(Luck, 2014). Note that the ex-Gaussian distribution is only a coarse approximation of a reaction time distribution (Matzke & Wagenmakers, 2009; Sternberg & Backus, 2015), but it has the advantage of simplicity and is sufficient for assessing ERP waveform distortions. Other families of functions could also be used to simulate ERP waves, such as the gamma distribution (Kummer et al., 2020).

4 More trials are typically used in experiments that examine smaller components, so an artifactual effect of 0.2 µV might be statistically significant in a typical N2pc experiment but would be unlikely to be significant in a typical P3 experiment.

